# Transcriptome-informed reduction of protein databases: an analysis of how and when proteogenomics enhances eukaryotic proteomics

**DOI:** 10.1101/2021.09.07.459229

**Authors:** Laura Fancello, Thomas Burger

## Abstract

**Background:** Proteogenomics aims to identify variant or unknown proteins in bottom-up proteomics, by searching transcriptome- or genome-derived custom protein databases. However, empirical observations reveal that these large proteogenomic databases produce lower-sensitivity peptide identifications. Various strategies have been proposed to avoid this, including the generation of reduced transcriptome-informed protein databases (*i.e.*, built from reference protein databases only retaining proteins whose transcripts are detected in the sample-matched transcriptome), which were found to increase peptide identification sensitivity. Here, we present a detailed evaluation of this approach.

**Results:** First, we established that the increased sensitivity in peptide identification is in fact a statistical artifact, directly resulting from the limited capability of target-decoy competition to accurately model incorrect target matches when using excessively small databases. As anti-conservative FDRs are likely to hamper the robustness of the resulting biological conclusions, we advocate for alternative FDR control methods that are less sensitive to database size. Nevertheless, reduced transcriptome-informed databases are useful, as they reduce the ambiguity of protein identifications, yielding fewer shared peptides. Furthermore, searching the reference database and subsequently filtering proteins whose transcripts are not expressed reduces protein identification ambiguity to a similar extent, but is more transparent and reproducible.

**Conclusion:** In summary, using transcriptome information is an interesting strategy that has not been promoted for the right reasons. While the increase in peptide identifications from searching reduced transcriptome-informed databases is an artifact caused by the use of an FDR control method unsuitable to excessively small databases, transcriptome information can reduce ambiguity of protein identifications.

## BACKGROUND

The term “proteogenomics” nowadays, indicates the combined analysis of genomics and/or transcriptomics with proteomics, hereby covering a broad spectrum of applications. Originally, it was adopted to refine genomic annotation by using peptide sequences identified by mass spectrometry as evidence for genomic coding regions^1^. While this is challenging for eukaryotes due to their larger genome size, efficient approaches were developed to better characterize the coding potential of prokaryotic genomes (1, 2). More recently, proteogenomics has been increasingly employed to discover small proteins, since short open reading frames typically are difficult to predict by conventional genome annotation tools (3, 4). Proteogenomic applications were also extended to the study of gene expression regulation at transcript and protein level or to the identification of cancer-specific protein variants (5, 6). Most importantly, proteogenomics emerged as an attractive strategy to enhance eukaryotic proteomics, which will be the focus of this work. Proteogenomics may enhance eukaryotic proteomics in two main ways^7^ : *i.* improving protein inference; *ii.* improving database searches for peptide identification.

Protein inference is a central issue in proteomics, given the presence of shared peptides, which are especially abundant in eukaryotes. These peptides might originate from different proteins sharing homology; or from distinct proteoforms due to alternative mRNA splicing, post-translational modifications, proteolytic cleavages, and/or allelic variants. Indeed, for experimental reasons, in bottom-up mass-spectrometry-based proteomics – the most widely used proteomic approach – peptide-protein connectivity is lost and protein identifications must be inferred from peptide identifications. Traditionally, the issue of protein inference was addressed using simple heuristics, such as the two-peptide rule (proteins must be identified by at least two peptides) or the parsimony principle (the smallest subset of proteins which can explain most or all peptides is retained) (7–9). Later, more refined approaches have been developed, which represent the mapping of shared peptide onto their parent proteins with probabilistic models (10–14) or which define different levels of peptide evidence based on gene models (*e.g.*, unambiguous, ambiguous from different isoforms or from different genes etc.) to infer a set of protein identifications with minimal false or ambiguous peptide assignments (15). Most commonly, when proteins cannot be distinguished based on peptide identifications (*i.e.*, they are identified by an identical set of peptides) they are reported as a protein group. This complicates comparisons between distinct experiments and protein quantification. In this context, proteogenomics can enhance protein inference using evidence from transcript expression: in particular, some Bayesian approaches have been developed based on this strategy (16–18). The other main contribution of proteogenomics to proteomics relates to refinements of the reference protein databases used for peptide identification. Classically, peptides produced by bottom-up proteomics are identified by matching their experimentally-measured mass spectra against theoretical spectra for all candidate peptides contained in a user-selected reference protein database. The underlying assumption is that the database exhaustively and accurately describes all the protein sequences present in the sample. However, this assumption may be unrealistic for two reasons. First, reference databases only contain canonical – experimentally-validated or predicted – protein sequences, but other variants or isoforms may be present, especially in tumor samples. Second, one may simply lack a reference protein database for less studied organisms for which scarce or no genomic annotation is available. In the first case, more exhaustive protein databases including undocumented or variant peptides can be generated by appending variant sequences from public genomic repositories (*e.g.*, COSMIC or dbSNP) to the reference database (19–21) or adding sample-specific variants identified from matched transcriptomes or genomes. In the second case, protein databases can be generated by 6-frame translation of the genome or sample-matched transcriptome (22, 23). A major downfall of these proteogenomic databases is that they tend to be very large with long and complex genomes such as for eukaryotes. Searching very large databases comes at a considerable computational cost, and complicates the task of discriminating between correct and incorrect matches. In particular, various studies have shown that when using target-decoy competition (TDC) for FDR control on large database searches, fewer peptides are identified at the same FDR level. This contrasts starkly with the initial reason for the development of proteogenomics (24, 25). To avoid reducing the number of identifications, it was proposed that FDR validation of canonical and novel peptides should be performed separately, and that post-search filters or machine-learning methods should be applied to increase confidence in the newly-identified peptides (19–21,26,27). In addition, various strategies were adopted to limit the size of databases generated by proteogenomics. When possible, 6-frame genome translation was replaced by translation of candidate ORFs identified by gene prediction algorithms or of *de novo* assembled RNA transcript sequences, when a sample matched transcriptome is available (28, 29). Alternatively, sample-specific variants from matched genome or transcriptome sequencing were preferentially added to the reference database rather than variant sequences from COSMIC or dbSNP (30–32). A more refined method using transcriptome information to refine protein inference in addition to peptide identification was also tested on prokaryotes (33). While only rarely used for these organisms, as the issue of database size is less important than for eukaryotes, it is suitable and particularly relevant for proteogenomics on eukaryotes. Finally, in some studies, after appending variants identified from sequencing data, the reference database was reduced to contain only proteins for which transcripts were detected by transcriptomics, since according to the “central dogma of biology”, there can be no protein without the corresponding transcript (34–36). Yet other groups proposed to generate reduced transcriptome-informed protein databases by barely reducing the reference database to proteins for which transcripts were detected, without including any novel sequences (34,36,37). The only declared objective of this approach is to increase the sensitivity of identification of known sequences. Indeed, the authors claim that searching such reduced transcriptome-informed databases can increase the number of valid identifications. A strategy was also proposed to optimize the balance between identifications lost due to the incompleteness of an excessively reduced database and additional identifications made from searches of reduced transcriptome-informed databases, to maximize the number of valid identifications (38).

However, in all these studies: *i.* Only limited attention was given to the mechanistic explanation for the increased number of peptides identified searching smaller databases; *ii.* Little is known about how reduced transcriptome-informed database searches affect protein inference in terms of ambiguity of protein identification and shared peptide assignments. Therefore, in this article, we investigated the use of reduced transcriptome-informed sample-specific protein databases, focusing on these two methodological aspects. Our investigations led to three conclusions. First, the reported increase in the number of identifications obtained by searching a reduced transcriptome-informed database is a statistical artifact. While the associated risk has previously been reported in a metaproteomics context (without proper mechanistic explanation (39, 40), we were able to establish that it is simply the spurious consequence of an underestimated FDR resulting from the sensitivity of TDC to database size (also reported in (41)). In other words, reducing the size of the database searched to increase sensitivity broadly amounts to validating peptide identifications at a higher-than-reported FDR. This observation consequently raises questions as to the validity of peptides that are only identified when searching the reduced database, and as to cross-study comparability. However, searching reduced transcriptome-informed protein databases followed by accurate FDR control remains nonetheless an interesting strategy, as it decreases the ambiguity of protein identifications by reducing the proportion of shared peptides and the size of protein groups. Finally, searches against the reference database followed by post-hoc filtering of proteins for which there is no evidence of transcript expression provides comparable proteomic identifications to searches against the reduced transcriptome-based database. This strategy guarantees better transparency and comparability between studies. To facilitate use of this search strategy, we produced an R package to perform post-hoc filtering on proteomic identifications based on transcript evidence that also allows manual inspection of protein identifications and shared peptide assignments within protein groups of interest while also providing information on transcript expression.

## RESULTS

### Reduced transcriptome-informed database search does not increase sensitivity if FDR is accurately controlled

To investigate how searching reduced transcriptome-informed protein databases affects proteomic identifications, we used four different samples (hereafter referred to as Jurkat, Lung, MouseColon and Spleen) for which matched transcriptome and proteome data were publicly available (**Additional File 1: Table S1**). For each of them, we built a sample-specific reduced transcriptome-informed protein database as a simple subset of the reference protein database, by including only those proteins whose corresponding transcript is expressed. Briefly, we first processed transcriptome datasets and identified the set of transcripts expressed in each sample using StringTie, a common transcriptome assembly method (see “Transcriptome analysis” in Methods). Then, we generated reduced databases for use in MS/MS searches by retaining from the Ensembl human protein database only proteins for which transcripts were expressed in the sample-matched transcriptome (see “Construction of reduced transcriptome-informed protein databases for MS/MS searches” in Methods) (Figure 1A-B). We compared valid peptide-spectrum matches (PSMs) obtained from the MS/MS search against the whole Ensembl human database (referred to as the “full database”) or against the sample-specific reduced database (referred to as the “reduced database”) at 1% FDR, as estimated by TDC. In agreement with previous studies (34,36,38), we found that a few spectra and peptides identified in the full database were lost when searches were performed against the reduced database (“lost in reduced DB”), whereas others were only identified in the reduced database search (“additional in reduced DB”) (Figure 1C). Lost identifications are due to incompleteness of the reduced database. Indeed, even more identifications were lost when using a further reduced protein database, such as that generated on the basis of the smaller set of expressed transcripts identified by Cufflinks, an alternative method of transcriptome assembly (**Additional File 2: Fig. S1**). Additional identifications, in contrast, are commonly ascribed to increased sensitivity of MS/MS searches when using smaller databases, such as reduced transcriptome-informed protein databases (34,36,38). In this study, we investigated the origin of these additional identifications more thoroughly. To do so, we performed a detailed comparison of all (target or decoy) PSMs retained from the full and reduced database searches, after applying validation prefilters (*i.e.*, single best-scoring PSM for each spectrum, minimum peptide length of 7 amino acids), but before filtering at 1% FDR (Figure 1A). Since the reduced database was constructed as a simple subset of the Ensembl human database and considered only a single best-scoring peptide for each spectrum (see “Proteome analysis” in Methods), we could easily map each spectrum match between the two searches. Two interesting observations emerged from this comparison.

**Figure 1.**
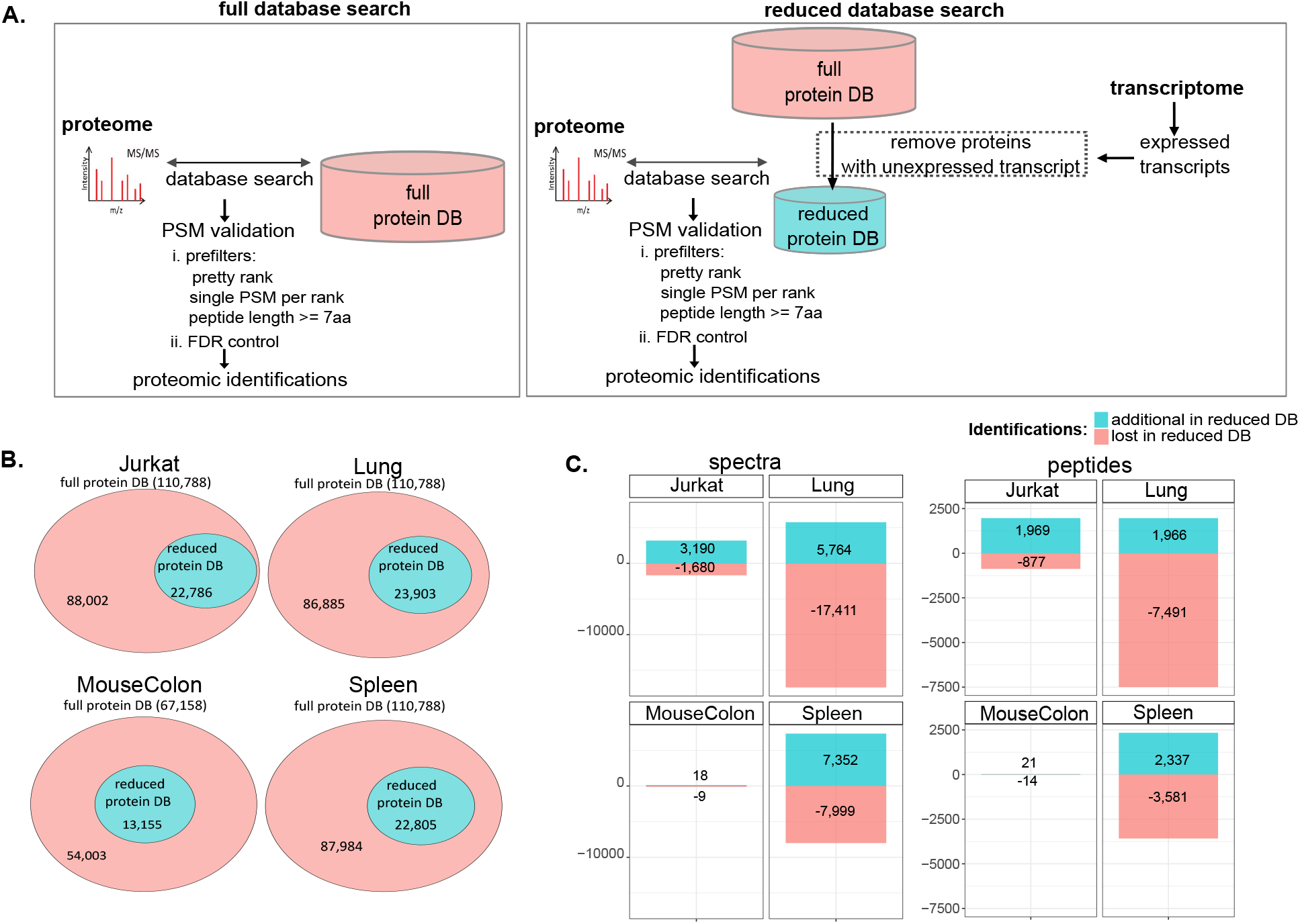
Comparison of proteomic identifications from the full reference database or the transcriptome-informed reduced database searches. **A.** Graphical representation of the two MS/MS search strategies compared here. MS/MS spectra were assigned searching either the reference human Ensembl protein database (full protein DB) or a subset of the reference database, generated based on transcript expression (reduced protein DB), using Mascot software. PSMs were first validated using Proline software with the following prefilters: *i.* PSMs with score difference < 0.1 were considered of equal score and assigned to the same rank (pretty rank); *ii.* only a single best-scoring PSM was retained per query (single PSM per rank); *iii.* minimum peptide length >= 7 amino acids. PSMs were then filtered at the score cutoff estimated by target-decoy competition for 1% FDR control. **B.** Size and overlap of the reference human Ensembl protein database (full protein DB) and the sample-specific reduced transcriptome-informed protein databases (reduced protein DB). **C.** Number of spectra (left) or peptides (right) exclusively identified in the reduced database (“additional in reduced DB”, in blue) or exclusively identified in the full database (“lost in reduced DB”, in red) searches.

First, several spectra were reallocated in the reduced database (*i.e.*, assigned to different matches in the reduced database and the full database). This phenomenon occurred when the peptide match from the full database was not present in the reduced database. However, in no case we observed a reallocation on a target with a higher score (Figure 2A and 2B**, Additional File 1: Table S2, Additional File 2: Fig. S2A and S2B**) This observation holds true independently of the search engine employed for peptide identification (**Additional File 1: Table S3**). Therefore, additional identifications in the reduced database do not come from an improved search score of target matches in the reduced database. For the sake of clarity, we emphasize that the few spectra which matched only sequences in the reduced database (indicated as “no match, target” or “no match, decoy” in Figure 2A**, Additional File 2: Fig. S2A, S3A, S4, S5, S6 and S7**) do not contradict this observation; they can all be explained by reallocation and the prefilters applied for validation (**Additional File 2: Fig. S3B,** see **Additional File 3: Supplementary Note 1** for a detailed explanation).

**Figure 2.**
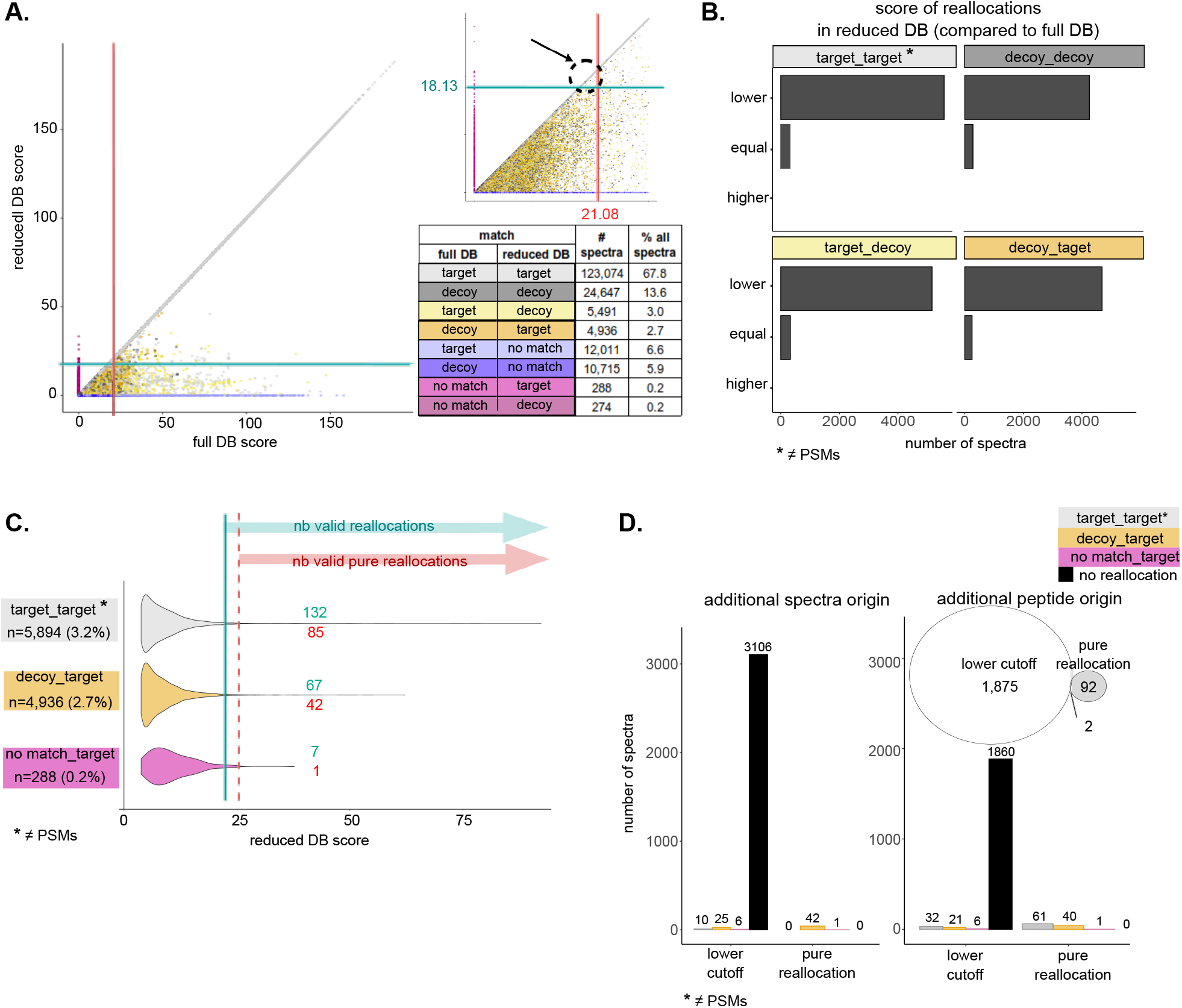
Lower cutoff for FDR control in the reduced database generates additional identifications (Jurkat). **A.** Scatter plot comparing for each spectrum its PSM score from the full (x axis) or reduced database (y axis) searches. A color code indicates the type of match (“target”, “decoy”, or “no match”) in the two searches. Score cutoffs obtained by TDC at 1% FDR are shown as red and blue lines for the full and reduced database, respectively. The upper-right insert zooms in on PSMs accepted at 1% FDR only in the reduced database, due to the lower score cutoff at 1% FDR (arrow pointing to the dashed circle). **B.** Number of reallocations whose score in the reduced database was equal to or lower or higher than the score in the full database. **C.** PSM scores for reallocations to target matches in the reduced database, grouped by the type of match in the full database. The number of reallocations passing the reduced database cutoff at 1% FDR is shown in blue (“nb valid reallocations”) and of those passing the full database cutoff at 1% FDR - additional valid identifications exclusively generated by reallocation, independent of the lower cutoff - in red (“nb valid pure reallocations”). **D.** Number of additional spectra (left) and number of spectra identifying additional peptides (right) exclusively identified in the reduced database search due to: *i.* lower score cutoff at 1% FDR in the reduced database compared to the full database – *i.e.*, PSMs only passing the cutoff from the reduced database search, including identical PSMs in both searches (black) and reallocations from target (gray), decoy (orange) or no match (magenta) in the full database to target matches in the reduced database; *ii.* pure reallocation – *i.e.*, additional identifications exclusively due to reallocation. The Venn diagram illustrates the corresponding non-redundant number of additional peptides (*i.e.*, not identified in the full database search) identified by these spectra.

The second main observation was that the score cutoff estimated by TDC at 1% FDR (*i.e.*, the score defining the set of accepted PSMs, while respecting the constraint that less than 1% of them are expected to be false discoveries) was lower for the reduced database than for the full database search (Figure 2A**, Additional File 2: Fig. S2A, Additional File 1: Table S4**). This was confirmed at various levels of FDR control (0.5%, 1%, and 5%) (and was also observed when using a different search engine than Mascot, namely MS-GF+ (**Additional File 1: Table S4** and **S5).** As a consequence of the lower score cutoff for FDR control estimated by TDC for the reduced database than for the full database, for a few spectra no match was validated after FDR control in the full database, whereas, at a lower or at most equal score, the match was validated in the reduced database search (pointed out by the arrow in Figure 2A and **Additional File 2: Fig. S2A**). This clearly explains why these PSMs are accounted for as additional identifications in the reduced database search. We also observed a few reallocations, which can likewise yield additional spectra and/or peptide identifications in the reduced database search. These cases are, in particular, reallocations from non-target matches in the full database to target matches in the reduced database search (up to 2.7% of all spectra, depending on the sample) and reallocations between different target matches (up to 3.6% of all spectra, depending on the sample) (Figure 2A,C **and Additional File 2: Fig. S2A,C, Additional File 1: Table S6**). However, only a minority of these reallocations are valid identifications when applying 1% FDR control (*i.e.*, pass the score cutoff for FDR control from the reduced database search) (Figure 2C**, Additional File 2: Fig. S2C, Additional File 1: Table S6**). Furthermore, even fewer of them would pass the cutoff determined for the full database search, and they are hereafter referred to as “pure reallocations” to indicate that additional identifications from these PSMs originate solely from reallocation and they do not represent additionally validated PSMs due to the lower cutoff for 1% FDR validation in the reduced database (Figure 2C**, Additional File 1: Table S5, Additional File 2: Fig. S2C and S8A**). Additional peptide identifications originating from either a lower cutoff for FDR control or from pure reallocations had lower PSM scores compared to peptides identified in both database searches (**Additional File 2: Fig. S8B**). For pure reallocations, the difference in score between the full and reduced database match was sometimes quite large, especially for target PSMs in the full database that were reallocated in the reduced database search (**Additional File 2: Fig. S8C**). Furthermore, additional peptide identifications only allowed between 6 and 8 supplementary protein identifications (*i.e.*, protein groups for which protein members were not identified in the full database search) per sample (**Additional File 1: Table S7**). Thus, these additional identifications are of lower quality and provide little benefit in terms of protein identification.

Overall, only a few additional identifications were obtained thanks to pure reallocations; all other additional identifications were obtained due to the lower cutoff for FDR control in the reduced database. Concretely, the lower cutoff explained between 98.8% and 94.4% of additional spectral identifications and between 95.2% and 77.5% of additional peptide identifications, depending on the sample (Figure 2D**, Additional File 2: Fig. S2D**, **Additional File 1: Table S8**). We next investigated the reasons why lower cutoffs were observed at a same FDR threshold in the reduced databases. To do so, we first simulated the outcome if the cutoff were instead equal to that of the full database (Figure 3A**, Additional File 2: Fig. S9A, Additional File 1: Table S9**). In these conditions, the proportion of valid decoys in the reduced database search was observed to considerably decrease compared to the full database, with a net loss of 27.1% to 50% valid decoys (Figure 3B, **Additional File 2: Fig. S9B, Additional File 1: Table S9**). Indeed, a significant fraction of spectra matching valid decoys in the full database were assigned to invalid or non-decoy matches in the reduced database, and were not counterbalanced by reallocations in the other direction (*i.e.*, from invalid/non-decoy matches to valid decoys) (Figure 3C**, Additional File 2: Fig. S9C, S4, S5, S6, S7 Additional File 1: Table S10**). In contrast, the majority of spectra matching valid targets in the full database matched the same valid target in the reduced database, so that their loss was quite limited (Figure 3C**, Additional File 2: Fig. S9C, Additional File 1: Table S10**). Although spectra matching valid decoys in the full database were reallocated much more frequently in the reduced database than spectra matching valid targets, upon reallocation they behaved similarly: few reallocations resulted in a valid match of the same type (Figure 3C, **Additional File 2: Fig. S9C, Additional File 1: Table S10**), and the difference in score between full and reduced database matches was comparable (**Additional File 2: Fig. S10A**). Hence, the proportion of valid decoys lost in the reduced database was higher than that of targets lost, simply because a higher proportion of decoys was reallocated. This observation can easily be explained by how the reduced database was generated: only proteins whose transcript is expressed, thus those more likely to be present, were retained from the canonical full protein database. Therefore, all valid targets from the full database are theoretically still present in the reduced database, but the same cannot be said for decoys, that by definition represent random matches (**Additional File 2: Fig. S10B**). The lower cutoff obtained by TDC for the reduced database allows a few more decoys to be validated and thus recovers the proportion of valid decoys required to declare a nominal FDR level of 1% (Figure 3B**, Additional File 2: Fig. S9B, Additional File 1: Table S9**). We suggest that the additional identifications validated at a lower cutoff in the reduced database are simply a byproduct of the known influence of database size on TDC (41), rather than evidence of increased sensitivity when searching reduced databases. Naturally, in the absence of a benchmark, it is impossible to determine whether these identifications represent correct matches that were missed in the full database due to FDR overestimation, or incorrect matches that were accepted in the reduced database due to FDR underestimation. Nevertheless, three main observations indicate that they should at least be considered with caution. First, these matches were accepted in the reduced database at quite low scores, meaning that, in any case, they represent low-quality spectra that cannot be identified with any great confidence. Second, it could be assumed that additional identifications arise due to the removal, in the reduced database, of high-scoring decoys that out-compete correct target matches, thus reducing sensitivity. However, in our study, most of the additional identifications did not represent reallocations from decoys to targets; rather, they consisted in the same PSMs that were accepted at 1% FDR only in the reduced database due to the lower score cutoff applied (Figure 2D-E**; Additional File 2: Fig. S2D-E, Additional File S1: Table S8**). Third, and most importantly, the artifactual origin of the additional identifications, given TDC sensitivity to database size, is supported by the comparison between the behavior of TDC and of the Benjamini-Hochberg (BH) procedure for FDR control (42). The BH procedure is known to yield a conservative and stable FDR control, and it was recently successfully applied to peptide identification (41). In particular, TDC was found to be less conservative and less stable than BH with respect to preliminary filters on precursor mass accuracy: at a narrower mass tolerance, fewer decoys were fair competitors for incorrect random matches, and consequently cutoffs were artificially lowered. Therefore, reducing the database size can similarly result in an insufficient number of decoys to accurately simulate incorrect target matches, thus leading to the observed lower cutoffs. To confirm this possibility, we applied the BH procedure to target-only database searches (see “Proteome analysis” in Methods). We obtained more conservative score cutoffs and, most importantly, more stable with respect to database size, compared to TDC and we confirmed this at various levels of FDR control (0.5%, 1%, 5%) (Figure 3D**, Additional File 2: Fig. S9D, S11A-B, Additional File S1: Table S11**). In line with this result, a much more limited number of additional identifications was validated in the reduced database searches when using the BH-based FDR control (**Additional File 2: Fig. S11C**). We also employed the BH procedure for FDR control on concatenated target-decoy database searches; while doing so is nonsensical from a practical data-processing viewpoint, from a statistical methodology viewpoint, it simplifies comparisons between BH and TDC stabilities. As expected, more conservative and stable score cutoffs were obtained with the BH procedure (**Additional File 2: Fig. S11A-B)**.

**Figure 3.**
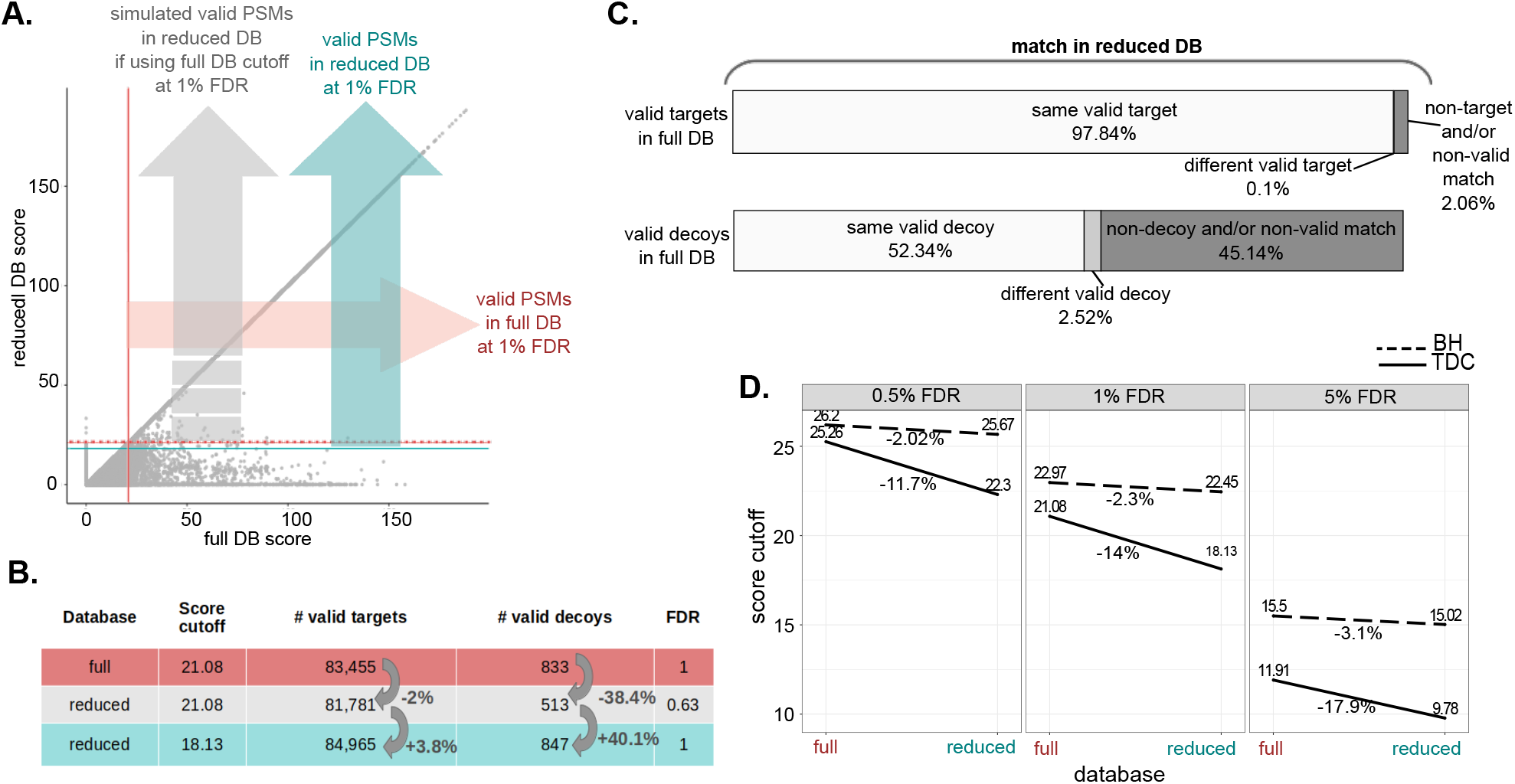
Lower cutoff for FDR control in the reduced database to recover valid decoys (Jurkat). **A.** Comparison of valid identifications obtained at 1% FDR from the full database (horizontal red arrow) or reduced database (vertical blue arrow) searches, and simulation of the valid identifications which would be obtained from the reduced database search if the score cutoff at 1% FDR were equal to that for the full database (dashed red arrow). **B.** Number of valid targets and decoys from the full or reduced database obtained at 1% FDR using the cutoffs estimated by TDC on the respective database search results (first and last rows). The second row presents the simulated number of valid targets and decoys which would be obtained from the reduced database if the estimated cutoff were the same as for the full database. Variations, expressed in percentages, are shown in gray. The associated nominal FDR level is reported (calculated as (*d*+1)/*t*, with *d* and *t* being the number of valid decoys and targets, as suggested in Levitsky *et al.* Proteome Res, 2016^46^. **C.** Match in the reduced database search for spectra matching valid targets or valid decoys in the full database. **D.** Score cutoffs obtained by TDC or by BH procedure for FDR control for the full or reduced database searches at various FDR levels (0.5%, 1%, and 5%). The variation in score cutoff between full and reduced database searches is reported as a percentage.

### Transcriptome information helps to reduce ambiguity of protein identifications

Although they do not enhance the sensitivity of peptide identifications, reduced transcriptome-informed databases can still be of benefit in proteomics at the protein-inference step, as they reduce the ambiguity of protein identifications. These databases include fewer proteins – only the proteins which are most likely to be present given their transcript expression – and it is reasonable to assume that with fewer possible protein matches, we might obtain fewer shared peptides and smaller protein groups. Such a decrease in protein group size has already been observed, but was either not discussed (43), or was attributed to an additional number of identifiable peptides available for parsimony-based protein inference (34). We have already demonstrated that additional identifications obtained when searching reduced databases actually derive from a flaw in TDC with respect to the reduced database size, and how they can be largely avoided by applying alternative FDR control procedures, such as BH. We will now show that straightforward searches against reduced databases followed by BH-based FDR control nonetheless yield smaller protein groups and less ambiguous protein identifications, thus regardless of additional identifications or protein inference methods.

Concretely, we compared identifications obtained from the full or reduced database searches followed by BH-based FDR control. The total number of identifications, at the spectrum, peptide, and protein level, was comparable (Figure 4A). For the number of protein-level identifications, we used the number of protein groups, as defined by the Proline software. Protein groups include the unambiguous identification of a single protein (single-protein groups) and groups of indiscernible proteins identified by the same sets of peptides (multi-protein groups) (see “Proteome analysis” in Methods). Interestingly, the proportion of single-protein groups was considerably higher for the reduced database (Figure 4B), meaning that protein identifications were less ambiguous.

**Figure 4.**
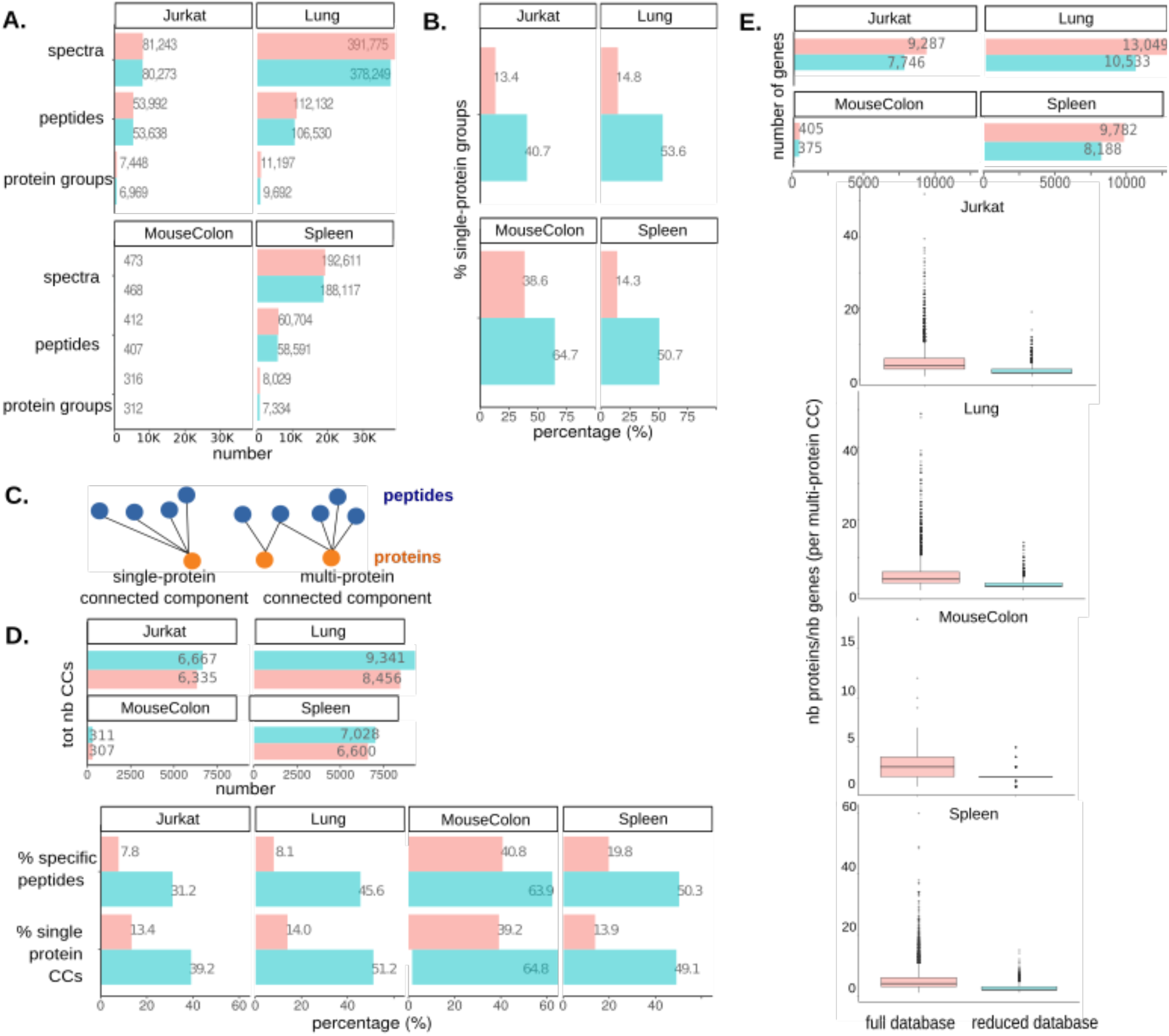
Transcriptome-informed reduced databases yield less ambiguous protein identifications. **A.** Number of valid identifications obtained from the full (red) or reduced (blue) target-only database searches, followed by BH procedure for 1% FDR control. The number of valid spectra, peptide and protein identifications are reported. Protein groups, as defined by the Proline software, represent protein identifications and include: *i.* proteins unambiguously identified by only specific peptides (*s*ingle-protein protein groups); *ii*. groups of proteins identified by the same set of shared peptides (multi-protein protein groups). **B.** Percentage of single-protein groups. **C.** Bipartite graph representation of peptide-to-protein mapping and exploitation of graph connected components to visualize and quantify the ambiguity of protein identifications. Unambiguous protein identifications are represented by CCs with a single protein vertex (single-protein CCs), while proteins sharing peptides are grouped in the same CC (multi-protein CCs) **D.** Upper panel: total number of connected components. Lower panel: percentage specific peptides and single-protein CCs. **E.** Genes encoding proteins from the full and reduced database searches. Upper panel: total number of genes associated with protein matches in the two searches. Lower panel: ratio between the number of protein members in each multi-protein CC and the number of genes encoding them.

We further characterized the ambiguity of protein identifications using the graph’s connected components. Briefly, we first represented peptide-to-protein mappings on bipartite graphs, with peptides and proteins as vertices and with edges featuring peptide-to-protein membership: this representation provides an easy picturing of the complex structures generated by shared peptides. Then, we calculated the connected components (CCs), *i.e.*, the largest subgraphs in which any two vertices are connected to each other by a path, and not connected to any of the other vertices in the supergraph. Proteins sharing one or more peptides are thus gathered in the same CC (multi-protein CCs), whereas unambiguous protein identifications are represented by CCs with a single protein vertex (single-protein CCs) (Figure 4C). As such, CCs constitute a peptide-centric strategy to represent ambiguous protein identifications and their shared peptides, and should not be confused with the classical protein-centric protein grouping strategy. We observed that, while the total number of CCs obtained was comparable, there was a considerably higher proportion of single-protein CCs in the graphs derived from the results of the reduced database search. Along with the reduction in protein group size, this is further evidence of decreased protein identification ambiguity (Figure 4D). Consistently, we also observed a greater proportion of specific peptides – and a correspondingly lower proportion of shared peptides – from the reduced database search (Figure 4D). Within multi-protein CCs, the ratio between the number of protein members and the corresponding number of their encoding genes was also lower for the reduced database, suggesting that at least part of the initial ambiguity occurred between proteins encoded by different genes (Figure 4E**, Additional File 1: Table S12**). In addition to a decrease in protein identification ambiguity, our observations indicated that searches against reduced databases were associated with lower ambiguity at the PSM level, although to a lesser extent (see **Additional File 3: Supplementary Note 2, Additional File 2: Fig. S12**). Finally, we adopted an alternative strategy to enhance proteomics by transcriptomics, which consists in an MS/MS search against the full database, followed by post-hoc filtering to remove proteins for which the corresponding transcript was not expressed and no specific peptide was identified (Figure 5A**, Additional File 2: Fig. S13**). The driving principle was to remove ambiguous protein identifications not supported by specific peptides or by transcriptomics, and thus to reduce ambiguity resulting from shared peptides. Intuitively, this approach can be pictured as less sensitive: some spectra could be filtered out (because of their matching sequence lacking transcriptomic evidence), instead of being possibly re-allocated onto the reduced database. However, knowing that pure reallocations are in practice scarce, we hypothesized that reduced database search or post-hoc filtering should yield similar results. This was empirically confirmed: Overall, we observed similar results to those obtained with reduced transcriptome-informed database searches and post-hoc filtering. First, a similar number of spectra and peptide identifications were obtained, comparable to that of the full database search (Figure 5B); second, these searches yielded a similarly increased proportion of single-protein CCs and specific peptides compared to full database searches (Figure 5C), indicating less ambiguous protein identifications. Post-hoc filtering is a transparent and easily interpretable approach, and we believe that it is highly suitable for use in studies seeking to enhance protein inference. While a few software tools already exist to generate reduced protein databases, we developed a specific toolbox of R scripts to perform the post-hoc filtering described. This toolbox also allows very efficient calculation of the CCs – which we have proposed as a means to quantify and compare ambiguity of protein identifications –, visualization of CCs of interest, and their manual inspection before and after post-hoc filtering.

**Figure 5.**
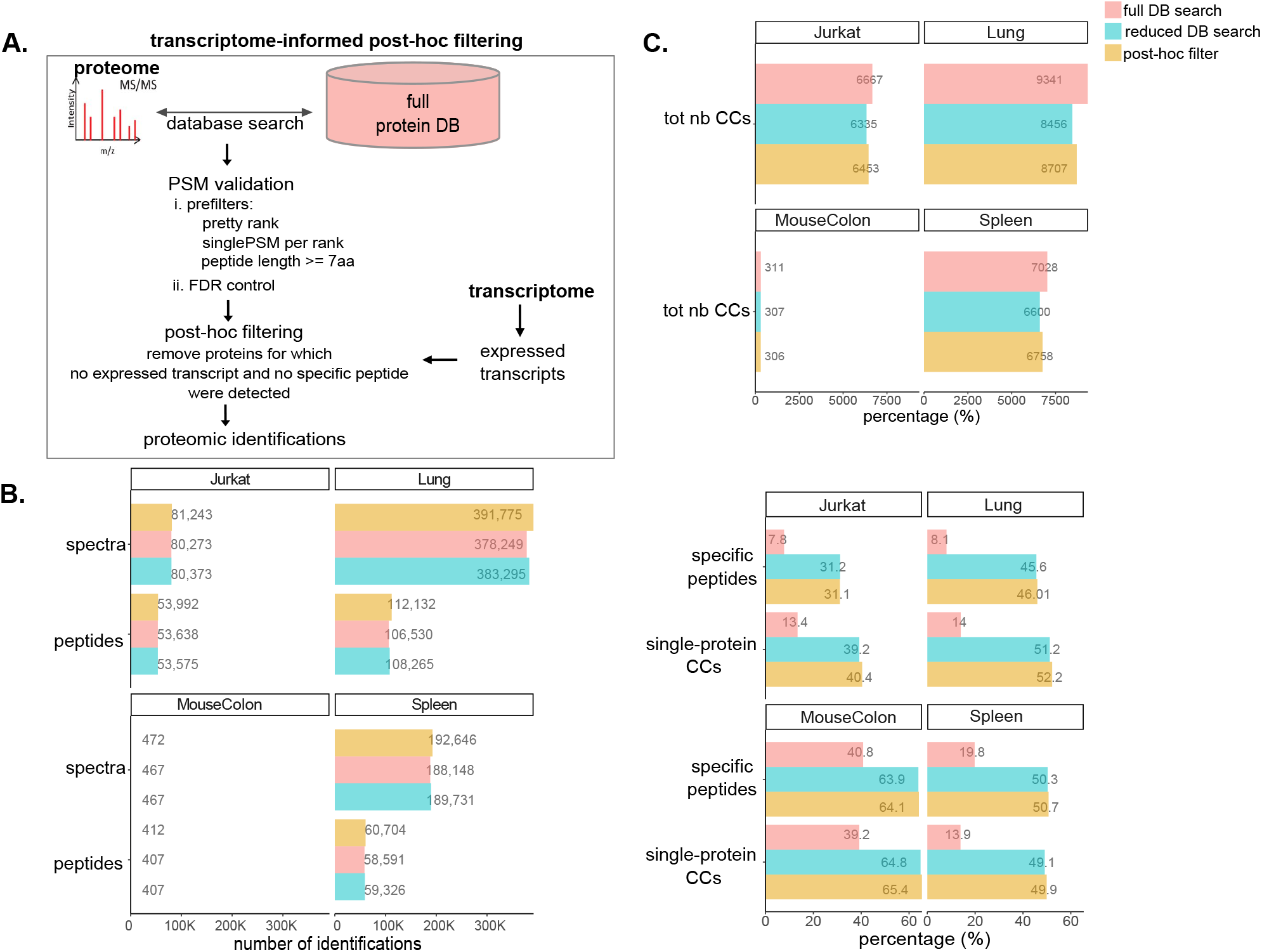
Transcriptome-informed post-hoc filtering and reduced database search strategies similarly reduce protein identification ambiguity. **A.** Illustration of the transcriptome-informed post-hoc filtering strategy. First, an MS/MS search was performed against the full canonical protein database. Then, proteins with no corresponding expressed transcript in the sample-matched transcriptome, and for which no specific peptide was detected (both conditions required) were removed. Peptides only mapping to that set of proteins were also removed. **B.** Number of valid spectra and peptide identifications obtained from the full or reduced target-only database search (red and blue) or from the post-hoc filtering strategy (orange), after 1% FDR control by BH procedure. **C.** Quantification of protein ambiguity for the full or reduced database search (red and blue) or the post-hoc filtering strategies (orange). Upper panel: total number of CCs obtained. Lower panel: percentage of specific peptides and single-protein CCs.

## DISCUSSION

In this article, we provide guidance for the mindful use of reduced transcriptome-informed protein databases for MS/MS searches. This type of reduced database stems from the attempt to counter excessive database inflation in proteogenomics studies on eukaryotes, when variant or novel proteoforms identified from sequencing data are added. Indeed, increased database size complicates the task of discriminating between correct and incorrect matches. When using TDC-based FDR control, inflated target databases come with an inflated number of decoys, and consequently a higher probability of high-scoring decoy matches. This has mainly been thought to reduce sensitivity of identifications in two ways. First, decoy matches may score better than correct target matches, thus out-competing them in spectrum-peptide assignment (“out-competing decoys”) and resulting in a decreased number of identifications. Second, decoys may have a higher probability of matching than incorrect targets, which violates the so-called Equal Chance Assumption of the TDC procedure and leads to an overestimated FDR, once again decreasing the number of identifications. As the main *raison d’être* of proteogenomics is to maximize the number of identifications, including variants or non-canonical peptides, much effort has been expended to avoid loss of sensitivity linked to excessively large databases, for example by reducing their size. While issues relating to the use of excessively large databases have been abundantly discussed, fewer authors have pointed out that excessively small databases may also be problematic, as they can also affect TDC estimations (41,44,45). With excessively small databases, TDC provides inaccurate FDR estimates, as they can only be asymptotically accurate (44,46,47). Furthermore, with too few (high-scoring) decoys, the probability of matching a decoy might be lower than the probability of matching an incorrect target, which once again violates the Equal Chance Assumption, but leads in this case to FDR underestimation and to an artifactual increase in identifications. In this study, we explicitly showed that the increased number of identifications obtained by searching reduced transcriptome-informed protein databases is most likely a statistical artifact due to the use of TDC on excessively small databases. We demonstrated how TDC estimates a lower score cutoff for 1% FDR control on the reduced databases compared to the full database search results. Consequently, some invalid PSMs in the full database will be retained as valid additional identifications only in the reduced database. We confirmed this observation at various levels of FDR control (0.5%, 1%, 5%) and for four different samples: three human-derived samples – healthy tissues (Lung and Spleen) or a cell line (Jurkat) – and one mouse-derived sample – flow cytometry sorted colon stem cells (MouseColon) –. The chosen samples represent different levels of proteomic complexity and come with a different number of spectra. Much fewer spectra were available for the MouseColon sample and slightly fewer for the Jurkat sample, which, interestingly, also presented a greater difference in score cutoffs between the full and reduced databases. Indeed, not only reduced database sizes but also a smaller number of spectra is believed to affect the ability of TDC to accurately estimate the FDR (44). We suggest that additional identifications obtained when searching such reduced databases are at least doubtful, and that it is unwise to employ reduced transcriptome-informed protein databases in an attempt to increase the number of identifications. Indeed, the additional identifications obtained had quite low scores and did not result from removal of out-competing decoys, a known cause of missed identifications in excessively large databases; rather, they represented identical PSMs in the two database searches but that were only accepted in the reduced database due to the lower score cutoff for the same level of FDR control. Most importantly, only a negligible number of additional identifications was generated from the reduced database search when using a method for FDR control that is known to be stable with respect to database size, such as BH. Indeed, using BH, the score cutoffs estimated for the full and reduced database searches, at the same level of FDR control, were almost identical. Thus, the BH procedure constitutes an interesting alternative to TDC for stable FDR control whatever the database size. However, many alternative approaches have been recently developed to cope with the weaknesses of classical TDC (48–53). It is important that proteogenomics researchers use one of these alternatives to avoid risking the propagation of statistical artifacts in their data. By doing so, they will no longer benefit from the hypothetical sensitivity increment assumed up to now, but this seems to be the necessary price to pay for rigorous control of the FDR

Reduced transcriptome-informed protein databases are nonetheless useful in bottom-up proteomics to reduce ambiguity in protein identifications due to the presence of shared peptides. In particular, we showed that searching a reduced database yielded a higher proportion of specific peptides and unambiguously identified proteins (*i.e*., single-protein CCs). Furthermore, the higher proportion of specific peptides and correspondingly lower proportion of shared peptides has a positive impact on precision in relative protein quantification. Indeed, when performing relative protein quantification, peptide abundances are used as a proxy for the abundance of their parent protein. In this scenario, shared peptides are difficult to handle, and since their relative abundance may depend on the contribution of multiple proteins, they are frequently discarded. This strategy has a downside, as it severely restricts the number of remaining quantifiable proteins – by reducing it to proteins with at least one specific peptide – and the amount of information available to estimate abundances – corresponding only to the number of specific peptides (54, 55). Therefore, a lower proportion of shared peptides represents more information available for quantification.

Finally, we showed that full database searches followed by post-hoc filtering of proteins for which no transcript is expressed provided comparable proteomic identifications to reduced database searches. This filtering approach similarly reduced ambiguity of protein identifications, while being more transparent and interpretable. We provide an R package (available on CRAN: https://CRAN.R-project.org/package=net4pg) to implement this type of post-hoc filtering strategy. The package allows to visualize ambiguous protein identifications and their peptides via bipartite graphs, to prune them based on transcript expression, and to manually inspect how this transcriptome-based post-hoc filtering strategy reduces ambiguity. Ambiguous protein identifications are represented and quantified using graph CCs, which constitute here subgraphs of proteins connected by shared peptides. This representation provides the following advantages: it is transparent, interpretable, non-redundant with respect to shared peptides, and independent of any strategy developed to define protein groups. Moreover, different options – which affect how shared peptides are treated – are available to the user to perform the transcriptome-informed filtering. The first option is to remove proteins with no transcript expression and then peptides only mapping on these proteins (either specific or shared by these proteins only). The second option is to remove proteins satisfying all the following criteria: no transcript expression and no specific peptides; then remove all peptides only shared between the removed proteins. The third option is to remove proteins satisfying all the following criteria: no transcript expression, no specific peptides and all their peptides being shared with at least one of the retained proteins, so that all peptide identifications are kept and only proteins are filtered out. While allowing a more pronounced reduction of ambiguity of protein identifications, we do not recommend the first option, which causes the reject of (even specific) peptide identifications and gives excessive weight to transcriptome information. Indeed, in some cases it can occur that a protein is expressed while its transcript is not detected (see below). By keeping proteins with no expressed transcript but a specific peptide identification, the other two filtering options better temper the weight of transcriptome information, while still providing less ambiguous protein identifications and therefore, more complete information for protein quantification. On the contrary, using reduced database searches, all proteins for which no transcript is detected are lost and this approach should thus be used with caution. Its underlying assumption that only proteins whose transcript is identified by transcriptomics are likely to be expressed may not hold true in some cases, due to, for example, differences between protein and mRNA half-lives or to the polyA enrichment protocol used for transcriptome generation (which misses the expression of non-polyA enriched transcripts, *e.g.*, histones, making the database incomplete). This drawback is shared with classic bottom-up proteomic approaches, where database exhaustiveness hampers sensitivity. To outstep this limitation, De Novo sequencing (14) has long been proposed, but as their performances are still limited, spectral rescoring has recently thrived as an interesting alternative (56). Results from this study are of interest also beyond proteogenomics. Indeed, database reduction is widely called for in proteomics, even though little attention is paid to the limitations of TDC when using excessively small databases. For example, some authors proposed to limit database size based on peptide detectability^30^, and it has been more generally claimed that “mass spectrometrists should only search for peptides they care about” (57). The statistical TDC artifacts observed with excessively small databases are an issue that also affects multi-step search strategies employed for conventional proteomics (58), for PTMs identification (59, 60) or for metaproteomics (39,61,62). The same issue might be encountered when searching peptide variants or modified peptides, which have higher false positive identification rates than canonical peptides or unmodified peptides (19,63,64). For this reason, it has been proposed to identify variant (respectively, modified) and canonical (respectively, unmodified) peptides using separate FDR validation (6,21,63,64). However, with separate searches or FDR validations, the variant database may become too small for accurate FDR validation of variant peptides, based on the conventional target-decoy approach and modifications of this approach are required (19, 63). Finally, beyond TDC limitations, the observation that transcriptome information can help decreasing ambiguity in protein identification is generally relevant in classical proteomics, but even more so in metaproteomics, which has to deal with an additional source of protein ambiguity: the presence of multiple organisms in the same sample.

## METHODS

### Proteogenomic datasets description

Four samples for which matched transcriptome and proteome were publicly available were analyzed: human healthy lung and spleen tissues (referred to as Lung and Spleen), a Jurkat cell line (referred to as Jurkat) and mouse colon stem cells (referred to as MouseColon). The lung and spleen samples were from a dataset by Wang *et al.* (*43*), which includes 29 histologically healthy human tissues generated to describe mRNA and protein expression levels throughout the human body. The Lung and Spleen transcriptome datasets were obtained by paired-end RNA sequencing on an Illumina HiSeq 2000/2500 system generating 2×100 base-long reads. The matching proteome datasets were obtained by quantitative label-free LC-MS/MS using an on-line nanoflow liquid chromatography system coupled to a Q Exactive Plus mass spectrometer, operating in data-dependent mode. Sample preparation included peptide fractionation via hSAX (hydrophilic strong anion) chromatography. Raw transcriptome and proteome data were downloaded from the EBI SRA (ArrayExpress accession: E-MTAB-2836; run accession: ERR315346) and the ProteomeXchange (dataset identifier: PXD010154; sample identifier: P013163) repositories, respectively.

The Jurkat cell line dataset was produced as part of a study by Sheynkman *et al.* (*31*). The Jurkat transcriptome dataset was obtained by paired-end RNA sequencing on an Illumina HiSeq 2000 system generating 2×200 bases long reads. The matched proteome dataset was obtained by quantitative label-free LC-MS/MS using a nanoAquity LC system chromatography system coupled to a Velos-Orbitrap mass spectrometer, operating in data-dependent mode. Sample preparation included peptide fractionation via high-pH LC separation. Transcriptome and proteome raw data were downloaded from NCBI’s Gene Expression Omnibus (GEO; accession: GSE45428) and the PeptideAtlas repositories (accession: PASS00215), respectively.

The mouse colon dataset comes from a study by Habowski *et al.* (*65*), which describes the transcriptome and proteome landscape of different types of epithelial colon cells, separated by flow cytometry. The transcriptome dataset was obtained by paired-end RNA sequencing on an Illumina HiSeq 4000 system generating 2×100 bases long reads. The matched proteome was obtained by quantitative label-free LC-MS/MS using an LC system chromatography system coupled to a Q Exactive Plus Orbitrap MS mass spectrometer, operating in data-dependent mode. Transcriptome and proteome data were downloaded from NCBI’s Gene Expression Omnibus (GEO; accession: GSE143915) and from ProteomeXchange (accession: PXD019351), respectively. Only one sample from the published dataset was analyzed, corresponding to the first replicate of colon stem cells and referred to as Stem BR#1 in the related publication.

### Transcriptome analysis

Raw reads were downloaded from public repositories and processed on the Galaxy platform available at https://usegalaxy.org/ (66) using common workflows for read preprocessing and alignment to identify expressed transcripts (**Additional File 1: Table S13**). First, sequencing adapters and low-quality (Phred score < 20) read ends were trimmed off using TrimGalore (https://www.bioinformatics.babraham.ac.uk/projects/trim_galore/), and reads shorter than 20 bp after trimming were discarded. Then, preprocessed reads were aligned against the corresponding reference genome (assembly GRCh38 for human samples and GRCm39 for the mouse sample) by the splice-aware STAR aligner (67) in default mode, using the Ensembl reference gene models for splice junctions. Only reads mapped to a proper pair, and passing platform quality controls were retained. Reads corresponding to optical or PCR duplicates were removed, as were non-primary and supplementary alignments. Initially, two common strategies for transcriptome assembly and quantification were applied: StringTie (68) and Cufflinks (69). Both programs were run to identify reference transcripts (and no novel transcripts), and two comparable sets of expressed transcripts were obtained. Unless otherwise specified, StringTie output was used for downstream analyses.

### Construction of reduced transcriptome-informed protein databases for MS/MS searches

For each sample, sample-specific protein databases were constructed for MS/MS search, containing only canonical sequences from the reference protein database for those proteins whose corresponding transcript is expressed in the sample. The obtained reduced database is therefore a simple subset of the protein reference database. Briefly, sample-matched transcriptomes were processed as described above, and the subsets of transcripts expressed at FPKM>1 were identified according to the StringTie or Cufflinks algorithms for transcript assembly and quantification. Then, the Ensembl reference protein database (GRCh38 and GRCm39 for human and mouse samples, respectively) was filtered to only retain proteins for which the corresponding transcript is expressed in the sample. For each sample, two sample-specific reduced versions of the Ensembl database were obtained, based on expressed transcripts from either StringTie or Cufflinks transcript quantification (**Additional File 2: Fig. S1A**). Unless otherwise specified, all downstream analyses were performed using the reduced transcriptome-informed database built according to expressed transcripts identified by StringTie, which is more recent than Cufflinks.

### Proteome analysis

Raw spectra were downloaded from public repositories and processed automatically using Mascot Distiller software (version 2.7, Matrix Science). PSMs were identified using Mascot search (version 2.6), if not otherwise specified. When specified, MS-GF+ (70) (version 2017.07.21) searches were performed on Galaxy (66) (https://usegalaxy.eu). Searches were performed against two different concatenated target-decoy databases: either the original Ensembl protein database (human GRCh38 and mouse GRCm39 for human and the mouse samples, respectively) or a reduced version of it containing only proteins for which the transcript is expressed (as described in the “Construction of reduced transcriptome-based protein databases for MS/MS searches” section). In both cases an equivalent number of decoy sequences was appended, as well as a custom database of common contaminant sequences (n=500) (and the corresponding number of decoys). Decoy sequences were generated by reversing target sequences running the perl script provided with Mascot software. Parameters used for Mascot (or MS-GF+) searches are reported in **Additional File 1: Table S14**.

The Proline software (71) was used for post-search PSM validation, applying the following prefilters: *i.* PSMs with score difference < 0.1 were considered of equal score and assigned to the same rank (pretty rank); *ii.* only a single best-scoring PSM was retained for each query (single PSM per rank); *iii.* minimum peptide length >= 7 amino acids. Prefiltered PSMs were then filtered at the score cutoff estimated for the 1% FDR control. Unless otherwise specified, the score cutoff for the FDR control was estimated by TDC (72). No protein inference was performed, but for each peptide, all possible protein matches were considered. Protein identifications were reported as protein groups, as defined in the Proline software. Protein groups included both the unambiguous identification of a single protein (single-protein groups) and groups of indistinguishable proteins identified by the same sets of peptides (multi-protein groups).

Further analyses were performed using the BH procedure for FDR control (42), as an alternative to TDC (see Results section “Transcriptome information helps to reduce ambiguity of protein identifications”). For these analyses, PSMs obtained from target-only protein databases, appended with the same database of common contaminant sequences were used, and searched, applying the same Mascot parameters as before.

### Bipartite peptide-protein graphs and connected components

Proteomic identifications were represented using bipartite graphs with two types of vertices – *i.* identified peptides; *ii.* all their possible proteins of origin – to more easily analyze and visualize groups of ambiguous protein identifications connected by shared peptides. Indeed, peptide assignments to proteins can generate very complex structures, when peptides are shared by different proteins, but they are easily represented using bipartite graphs. CCs from the graph – defined as the largest subgraphs in which any two vertices are connected to each other by a path and not connected to any other of the vertices in the supergraph – were then used to quantify the level of ambiguity of protein identifications.

To build bipartite graphs of proteomic identifications, first a tab-separated file containing identified peptides and all proteins they map on (one protein per line) was generated based on the output of PSM validation by Proline software. This file was then converted into an incidence matrix, with proteins along the columns and peptides along the rows, using the crosstab function from the GNU datamash program (http://www.gnu.org/software/datamash). The cross-product of the incidence matrix was used to produce the corresponding adjacency matrix, which describes protein-to-protein connections, based on shared peptides. Finally, CCs were calculated using the connComp() function from the “graph” R package applied to the adjacency matrix. Two types of CCs were defined: *i.* those containing a single protein (single-protein CCs), with only specific peptides, which constitute unambiguous protein identifications; *ii.* those containing multiple proteins sharing peptides (multi-protein CCs), which represent ambiguous protein identifications. Ambiguous protein identifications can be visually inspected by taking the CC of interest, extracting all specific and shared peptides mapping to the CC protein members from the incidence matrix and plotting peptide-to-protein mappings as bipartite graphs, using the “*igraph”* R package.

To decrease the computational cost when dealing with very large datasets or if computational resources are scarce, an alternative strategy was also developed to calculate CCs (**Additional File 2: Fig. S14**). First, the incidence matrix was reduced by removing all proteins not sharing peptides and all peptides unique to these proteins. Then, the corresponding adjacency matrix was generated as the cross-product of the incidence matrix. CCs can be more rapidly calculated on this reduced adjacency matrix. In this case, only multi-protein CCs are obtained, since protein identifications with only specific peptides, corresponding to single-protein CCs, were removed from the incidence matrix. While multi-protein CCs are those of interest when investigating ambiguous protein identifications from shared peptides, single-protein CCs can still be easily retrieved from the original incidence matrix if necessary.

A companion R code is provided as an R package (available on CRAN: https://CRAN.R-project.org/package=net4pg) implementing all the above-described steps, including: *i.* generating the adjacency matrix; *iii.* calculating CCs; *iii.* visualizing CCs as bipartite graphs.

### Transcriptome-informed post-hoc filtering

As an alternative to searching a reduced transcriptome-informed database, a transcriptome-informed post-hoc filtering strategy was tested. First, peptide identifications were obtained by searching the full reference protein database, and validated using Proline software, as described in the “Proteome analysis” section. An incidence matrix was generated to encode peptide-to-protein mappings (see the “Bipartite peptide-protein graphs and connected components” section). Then, the sample-matched transcriptome was analyzed to identify the set of transcripts expressed. Finally, transcriptome-informed filtering was performed using an in-house R code now available on CRAN as an R package (https://CRAN.R-project.org/package=net4pg). Three different options are available in the package to perform the filtering. The first option is to remove proteins with no transcript expression and peptides only mapping on these proteins (either specific or shared by these proteins only). The second option is to remove proteins satisfying all the following criteria: no transcript expression and no specific peptides; then remove all peptides only shared between the removed proteins. The third option is to remove proteins satisfying all the following criteria: no transcript expression, no specific peptides and all their peptides being shared with at least one of the retained proteins, so that all peptide identifications are kept and only proteins are filtered out. In this work, we employed the second option: the peptide-to-protein incidence matrix was filtered by removing proteins for which no expressed transcript and no specific peptide were detected. The one-to-one transcript-to-protein correspondence is guaranteed by the use of Ensembl as the reference protein database in proteomics and for genome annotation in transcriptomics. The filtered incidence matrix was then converted to an adjacency matrix to calculate CCs, as previously described (see “Bipartite peptide-protein graphs and connected components” section).

For this study, the post-hoc filtering strategy on PSMs obtained from searching the target-only full Ensembl protein database was used, followed by the BH procedure for FDR control, to allow comparison with the approach searching target-only reduced transcriptome-informed protein databases, followed by the BH procedure for FDR control. Indeed, the BH procedure was used as an alternative to TDC after searching reduced transcriptome-informed protein databases to obtain accurate FDR control, as explained in the Results section (see “Reduced transcriptome-informed database search does not increase sensitivity if FDR is accurately controlled”). However, in other contexts, post-hoc filtering could also be performed using PSMs from concatenated target-decoy searches followed by TDC-based FDR control.

## ABBREVIATIONS

BH: Benjamini-Hochberg procedure
CC: connected component
DB: database
FDR: false discovery rate
MS/MS: tandem mass spectrometry
PSM: peptide-spectrum match
TDC: target-decoy competition

## DECLARATIONS

### Ethics approval and consent to participate

Not applicable

### Consent for publication

Not applicable

### Availability of data and materials

All datasets analyzed in this study correspond to publicly-available data from the works of Wang et al.^33^ Sheynkman et al.^24^ and Habowski et al.^65^.

The transcriptome datasets were downloaded from the EBI SRA with ArrayExpress accession: E-MTAB-2836 (run accession: ERR315346), and from the NCBI Gene Expression Omnibus (GEO) with accession GSE45428. The proteome datasets were downloaded from PeptideAtlas with accession PASS00215 and from the ProteomeXchange repository with dataset identifier: PXD010154 (sample identifier: P013163).

Scripts used to perform these analyses are available as a platform-independent R package, under GPL-3 license, on the CRAN repository (https://CRAN.R-project.org/package=net4pg). A detailed description of the package usage is also available as a vignette on CRAN.

### Competing interests

The authors have no competing interests to declare.

### Funding

This work was supported by grants from the French National Research Agency: ProFI project (ANR-10-INBS-08), GRAL project (ANR-10-LABX49-01), DATA@UGA and SYMER projects (ANR-15-IDEX-02) and MIAI @ Grenoble Alpes (ANR-19-P3IA-0003).

### Author’s contributions

LF collected and reprocessed the data, developed the R package and drafted the manuscript. TB devised the experimental design and proposed algorithmic solutions. Both authors jointly interpreted the results and finalized the manuscript, which they approve.

## Acknowledgments

The authors are grateful to the IFB (Institut Français Bioinformatique, ANR-11-INBS-0013) infrastructure for providing computational resources.

## Notes

https://github.com/laurafancello/net4pg

### Competing Interest Statement

The authors have declared no competing interest.

