## Additional File 2 for "Transcriptome-informed reduction of protein databases: an analysis of how and when proteogenomics enhances eukaryotic proteomics"

### SUPPLEMENTARY FIGURE LEGENDS

**Figure S1. Two different methods for transcript assembly and their effect on reduced transcriptome-informed databases for proteomics. A.** Size (*i.e.*, number of protein sequences) and overlap between the full Ensembl human protein database ("full protein DB") and two reduced transcriptome-informed protein databases generated from it ("Cufflinks reduced DB" and "StringTie reduced DB"). The two reduced databases are based on slightly different sets of expressed transcripts, as identified in sample-matched transcriptome by two common transcriptome assembly methods: Cufflinks and StringTie. **B.** Additional or lost identifications in reduced database compared to the full database search. The reduced database based on Cufflinks transcriptome assembly is associated with a higher number of lost identifications than the one based on StringTie, because it is smaller (*i.e.,* higher incompleteness). All further analyses shown in this study are based on StringTie transcriptome assembly (see "Construction of reduced transcriptome-informed protein databases for MS/MS searches" in Methods).

**Figure S2. Lower cutoff for FDR control in the reduced database generates additional identifications (Lung). A.** Scatter plot comparing for each spectrum its PSM score from the full (x axis) or reduced database (y axis) searches. A color code indicates the type of match ("target", "decoy", or "no match") in the two searches. Score cutoffs obtained by TDC at 1% FDR are shown as red and blue lines for the full and reduced database, respectively. The upper-right insert zooms in on PSMs accepted at 1% FDR only in the reduced database, due to the lower score cutoff at 1% FDR (arrow pointing to the dashed circle). **B.** Number of reallocations whose score in the reduced database was equal to or lower (never higher) than the score in the full database. **C.** PSM scores for reallocations to target matches in the reduced database, grouped by the type of match in the full database. The number of reallocations passing the reduced database cutoff at 1% FDR is shown in blue ("nb valid reallocations") and of those passing the full database cutoff at 1% FDR - additional valid identifications exclusively generated by reallocation, independent of the lower cutoff - in red ("nb valid pure reallocations"). **D.** Number of additional spectra (left) and number of spectra identifying additional peptides (right) exclusively identified in the reduced database search due to: *i.* lower score cutoff at 1% FDR in the reduced database compared to the full database – *i.e.*, PSMs only passing the cutoff from the reduced database search, including identical PSMs in both searches (black) and reallocations from target (gray), decoy (orange) or no match (magenta) in the full database to target matches in the reduced database; *ii.* pure reallocation – *i.e.*, additional identifications exclusively due to reallocation. The Venn diagram illustrates the corresponding non-redundant number of additional peptides (*i.e.*, not identified in the full database search) identified by these spectra.

**Figure S3. Same spectrum PSM score in reduced database is never higher than in full database search. A.** Spectra with a match only in the reduced database (“no match, target”, “no match, decoy”) are represented with arbitrary score of 0 in the full database. In fact, these spectra have no match in the full database search, therefore no score either. **B.** Illustration of why some spectra may have no match in the full database, while having a match in the reduced database search. In all cases the rank 1 PSM in the full database did not pass the peptide length prefilter (>= 7 aa) and was not retained; such peptide being not present in the reduced database, the rank 1 PSM in the reduced database is another peptide, which satisfies the length prefilter. **C.** The combined use of “pretty rank” and “single PSM per rank” in PSM validation may cause some spectra to have a PSM score in the reduced database apparently higher than in the full database. **D.** Illustration of how the combined use of “pretty rank” and “single PSM per rank” in PSM validation may cause some spectra to have a PSM score in the reduced database apparently higher than in the full database. “Pretty rank” assigns the same rank to all PSMs from the same spectrum differing in score of less than 0.1. This may result in multiple PSMs of rank 1, with a slight difference in score (up to 0.1). The “single PSM per rank” filter retains a single best PSM per spectrum and, in some cases, the rank 1 PSM with the lower score is retained for the full database, while the higher scoring one is retained in the reduced database.

**Figure S4. Detailed view of PSMs obtained from the full or reduced database searches, split by their match type in each database search (Jurkat).** Each data point represents a spectrum, whose PSM score in the full and reduced database searches is reported by the x and y coordinates, respectively. Spectra are represented grouped by the type of match (target, decoy or no match) in each of the two searches (*<match fullDB>_<match reduced DB>*), which is also indicated by the color code. Score cutoffs estimated by TDC for 1% FDR control on the full and reduced database searches are represented as solid red and blue lines, respectively, on both the x and y axes. We also show on the y axis (representing reduced database search scores), the score cutoff estimated by TDC for the full databases search (dashed red line), to simulate what would occur if full and reduced database searches had the same cutoff for 1% FDR control. In each plot section delimited by these cutoffs, the corresponding number and percentage of PSMs are reported. Red circles highlight valid decoys in the full database search, which would be lost in the reduced database search if the score cutoff were the same as that from the full database (dashed red line). Green circles highlight valid decoys recovered in the reduced database search using the lower cutoff obtained by TDC for the reduced database (blue line).

**Figure S5. Detailed view of PSMs obtained from the full or reduced database searches, split by their match type in each database search (Lung).** Each data point represents a spectrum, whose PSM score in the full and reduced database searches is reported by the x and y coordinates, respectively. Spectra are represented grouped by the type of match (target, decoy or no match) in each of the two searches (“<match fullDB>_<match reduced DB>”), which is also indicated by the color code. Score cutoffs estimated by TDC for 1% FDR control on the full and reduced database searches are represented as solid red and blue lines, respectively, on both the x and y axes. We also show on the y axis( representing reduced database search scores) the score cutoff estimated by TDC for the full databases search (dashed red line), to simulate what would happen if full and reduced database searches had the same cutoff for 1% FDR control. In each plot section delimited by these cutoffs, the corresponding number and percentage of PSMs are reported. Red circles highlight valid decoys in the full database search, which would be lost in the reduced database search if the score cutoff were the same as that from the full database. Green circles highlight valid decoys recovered in the reduced database search using the lower cutoff obtained by TDC for the reduced database.

**Figure S6. Detailed view of PSMs obtained from the full or reduced database searches, split by their match type in each database search (MouseColon).** Each data point represents a spectrum, whose PSM score in the full and reduced database searches is reported by the x and y coordinates, respectively. Spectra are represented grouped by the type of match (target, decoy or no match) in each of the two searches (“<match fullDB>_<match reduced DB>”), which is also indicated by the color code. Score cutoffs estimated by TDC for 1% FDR control on the full and reduced database searches are represented as solid red and blue lines, respectively, on both the x and y axes. We also show on the y axis( representing reduced database search scores) the score cutoff estimated by TDC for the full databases search (dashed red line), to simulate what would happen if full and reduced database searches had the same cutoff for 1% FDR control. In each plot section delimited by these cutoffs, the corresponding number and percentage of PSMs are reported. Red circles highlight valid decoys in the full database search, which would be lost in the reduced database search if the score cutoff were the same as that from the full database. Green circles highlight valid decoys recovered in the reduced database search using the lower cutoff obtained by TDC for the reduced database.

**Figure S7. Detailed view of PSMs obtained from the full or reduced database searches, split by their match type in each database search (Spleen).** Each data point represents a spectrum, whose PSM score in the full and reduced database searches is reported by the x and y coordinates, respectively. Spectra are represented grouped by the type of match (target, decoy or no match) in each of the two searches ("<match fullDB>_<match reduced DB>"), which is also indicated by the color code. Score cutoffs estimated by TDC for 1% FDR control on the full and reduced database searches are represented as solid red and blue lines, respectively, on both the x and y axes. We also show on the y axis( representing reduced database search scores) the score cutoff estimated by TDC for the full databases search (dashed red line), to simulate what would happen if full and reduced database searches had the same cutoff for 1% FDR control. In each plot section delimited by these cutoffs, the corresponding number and percentage of PSMs are reported. Red circles highlight valid decoys in the full database search, which would be lost in the reduced database search if the score cutoff were the same as that from the full database. Green circles highlight valid decoys recovered in the reduced database search using the lower cutoff obtained by TDC for the reduced database.

**Figure S8. Additional peptide identifications. A.** Venn diagram illustrating the sources of additional identifications in reduced database searches (compared to full database): *i.* pure reallocation; *ii.* lower score cutoff estimated by TDC for 1% FDR control in the reduced database search (compared to the full database). Additional identifications originated by the lower cutoff can include reallocated spectra (intersection between lower cutoff set and reallocation set) and non-reallocated spectra ("lower cutoff" set, excluding the intersecting area). Pure reallocations are defined as those reallocations which would be validated even in the case where the score cutoff for 1% FDR control on the reduced database search results were the same as the full database cutoff ("reallocation" set, excluding the intersecting area). **B.** Distribution of PSM scores in the reduced database search for additional peptides exclusively identified in the reduced database search by pure reallocation or lower cutoff ("additional peptides in reduced DB"), and for peptides identified in both full and reduced database searches ("other peptides"). Top panel: Jurkat; bottom panel: Lung. **C.** Distribution of the score difference between the full and reduced database PSM for the same spectrum ("score difference, full DB score – reduced DB score"). Only pure reallocations which allow to identify additional peptides (compared to the full database) are shown. Pure reallocations are grouped according to the match type in the full database ("<match in full DB>_<match in reduced DB>"). Top panel: Jurkat; bottom panel: Lung.

**Figure S9. Lower cutoff for FDR control in the reduced database to recover valid decoys (Lung). A.** Comparison of valid identifications obtained at 1% FDR from the full database (horizontal red arrow) or reduced database (vertical blue arrow) searches, and simulation of the valid identifications which would be obtained from the reduced database search if the score cutoff at 1% FDR were equal to that for the full database (dashed red arrow). **B.** Number of valid targets and decoys from the full or reduced database obtained at 1% FDR using the cutoffs estimated by TDC on the respective database search results (first and last rows). The second row presents the simulated number of valid targets and decoys which would be obtained from the reduced database if the estimated cutoff were the same as for the full database. Variations, expressed in percentages, are shown in gray. The associated nominal FDR level is reported (calculated as $(d+1)/t$, with $d$ and $t$ being the number of valid decoys and targets, as suggested in Levitsky *et al.* Proteome Res, 2016[36]. **C.** Match in the reduced database search for spectra matching valid targets or valid decoys at 1% FDR in the full database. **D.** Score cutoffs obtained by TDC or by BH procedure for FDR control for the full or reduced database searches at various FDR levels (0.5%, 1%, and 5%). The variation in score cutoff between full and reduced database searches is reported as a percentage.

**Figure S10. Target and decoy reallocations in the reduced database search. A.** Normalized distribution of the difference in score between full and reduced database matches, for the same spectrum, upon reallocation. All (valid or invalid at 1% FDR) spectra reallocated between the full and reduced database searches are shown. Data are grouped according to the reallocation type, which is the type of match in the full and reduced database searches (target_target, decoy_decoy, target_decoy, decoy_target). **B.** Illustration of why a higher proportion of valid decoy rather than valid target matches in the full database is lost in the reduced database search, if the estimated cutoff for FDR control were the same as for the full database. The reduced database (in blue) is generated as a subset of the full database (in red) which only contains proteins whose transcript is expressed and thus more likely to be present. Therefore, all valid targets from the full database (indicated by a capital "T") theoretically are still present in the reduced database while this is not the case for valid decoys (indicated by a capital "D"), which, by definition, represent random hits.

**Figure S11. PSMs obtained from the full or reduced database searches, followed by BH procedure for FDR control**. **A.** Scatter plot comparing PSMs obtained searching the full or reduced target-only database searches, passing prefilters but prior to FDR control. Each data point represents a spectrum: its corresponding PSM score in the full and reduced database searches is reported on the x and y coordinates, respectively. A color code is used to represent the type of match ("target", or "no match") for each spectrum in the two searches. Score cutoffs obtained by BH at 1% FDR are also shown as red and blue lines for the full and reduced database searches, respectively. **B.** Score cutoffs estimated for the full or reduced databases at 0.5%, 1% or 5% FDR, using the TDC or BH method for FDR control. Three approaches were compared: *i*. concatenated target-decoy searches followed by TDC for FDR control; *ii*. target-only searches followed by BH procedure for FDR control; *iii*. concatenated target-decoy searches followed by BH procedure for FDR control, to provide a fairer comparison with TDC. The variation of score cutoff between full and reduced database searches is reported in percentage. **C.** Number of spectra (on the left) or peptides (on the right) exclusively identified in the reduced database ("additional in reduced DB" in blue) or exclusively identified in the full database ("lost in reduced DB" in red) search, using TDC or BH procedure for 1% FDR control on concatenated target-decoy database search results. The net difference between additional and lost identifications in the reduced database is also reported on top of each bar ("net").

**Figure S12. Searching reduced databases yields fewer rank 1 PSMs per spectrum.** **A.** Proportion of spectra with one or more equally good best matches ("# rank 1 matches per spectrum: 1 or >1”). **B.** Frequency distribution of spectra with 2 to *n* valid PSMs of rank 1.

**Figure S13. Transcriptome-informed post-hoc filtering strategy.** Illustration of the transcriptome-informed post-hoc filtering strategy. The upper graph represents peptide-to-protein mappings for valid proteomic identifications obtained from searching the reference protein database (full protein database). The graph is pruned by removing proteins with no expression of the corresponding transcript and no specific peptides (protein 3 in the example). All peptides exclusively mapping to these proteins are also removed.

**Figure S14. Illustration of the strategy to calculate efficiently the set of connected components on large bipartite graphs**. Peptide-to-protein mappings obtained from database search and PSM validation steps are reported as an incidence matrix with peptides as rows and proteins as columns. Each cell indicates whether the corresponding peptide maps on that protein (1) or not (0). All specific peptides and all those proteins which are only identified by specific peptides are removed from the incidence matrix, generating a reduced incidence matrix. The cross-product of the reduced incidence matrix is used to generate an adjacency matrix describing protein-to-protein connections, *i.e.*, whether two protein are identified by at least one shared peptide (1) or not (0). Same-protein connections on the diagonal are removed because of no interest (“remove self-loops”). Connected components are calculated from the adjacency matrix: they represent sets of proteins sharing peptides and are employed to visualize and quantify ambiguity of protein identifications. Each connected component of interest can then be visualized as a bipartite graph by recovering all specific and shared peptides mapping on its protein members from the original incidence matrix.

**Supp. Fig. 1**

**A.**

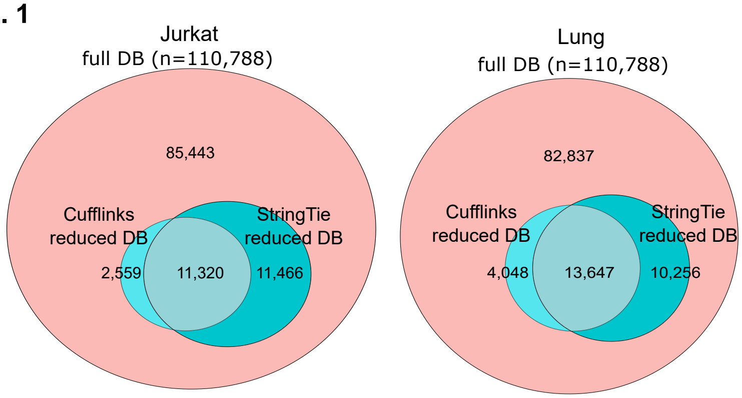

Jurkat
full DB (n=110,788)
85,443
Cufflinks reduced DB
StringTie reduced DB
2,559
11,320
11,466
Lung
full DB (n=110,788)
82,837
Cufflinks reduced DB
StringTie reduced DB
4,048
13,647
10,256


**B.**

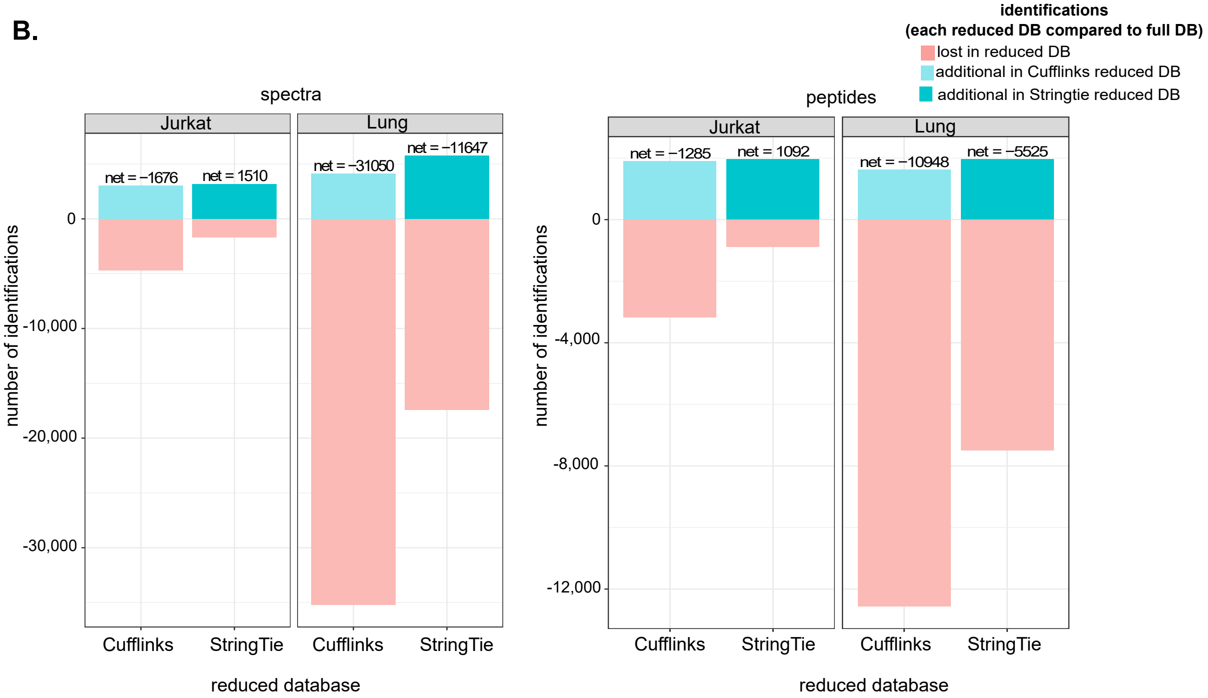

identifications
(each reduced DB compared to full DB)
lost in reduced DB
additional in Cufflinks reduced DB
additional in Stringtie reduced DB
spectra
Jurkat
Lung
net = −1676
net = 1510
net = −31050
net = −11647
0
-10,000
-20,000
-30,000
number of identifications
Cufflinks
StringTie
Cufflinks
StringTie
reduced database
peptides
Jurkat
Lung
net = −1285
net = 1092
net = −10948
net = −5525
0
-4,000
-8,000
-12,000
number of identifications
Cufflinks
StringTie
Cufflinks
StringTie
reduced database

#### Supp. Fig. 2

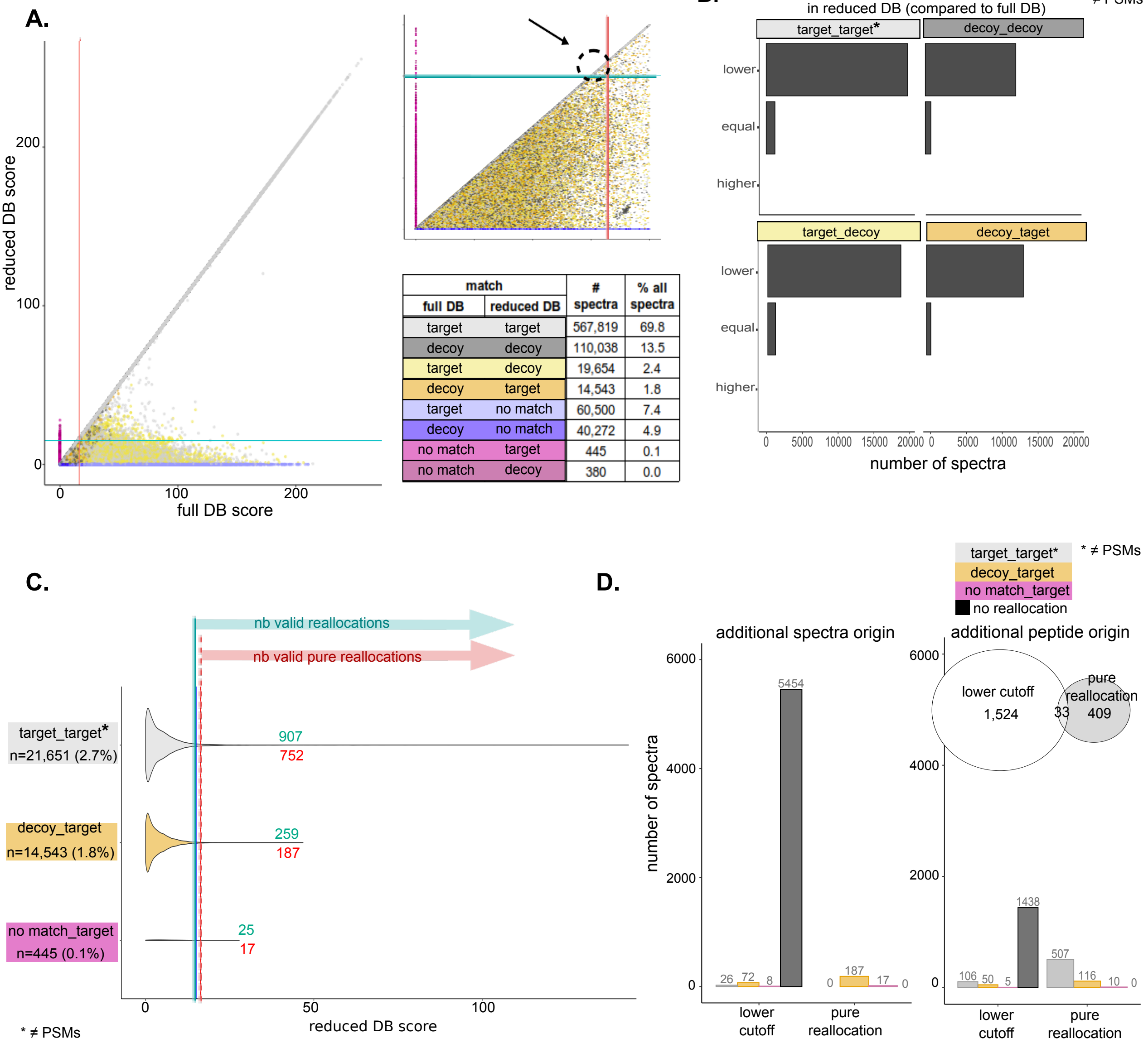


| match | | # spectra | % all spectra |
|---|---|---|---|
| full DB | reduced DB | | |
| target | target | 567,819 | 69.8 |
| decoy | decoy | 110,038 | 13.5 |
| target | decoy | 19,654 | 2.4 |
| decoy | target | 14,543 | 1.8 |
| target | no match | 60,500 | 7.4 |
| decoy | no match | 40,272 | 4.9 |
| no match | target | 445 | 0.1 |
| no match | decoy | 380 | 0.0 |

Supp. Fig. 3

A.

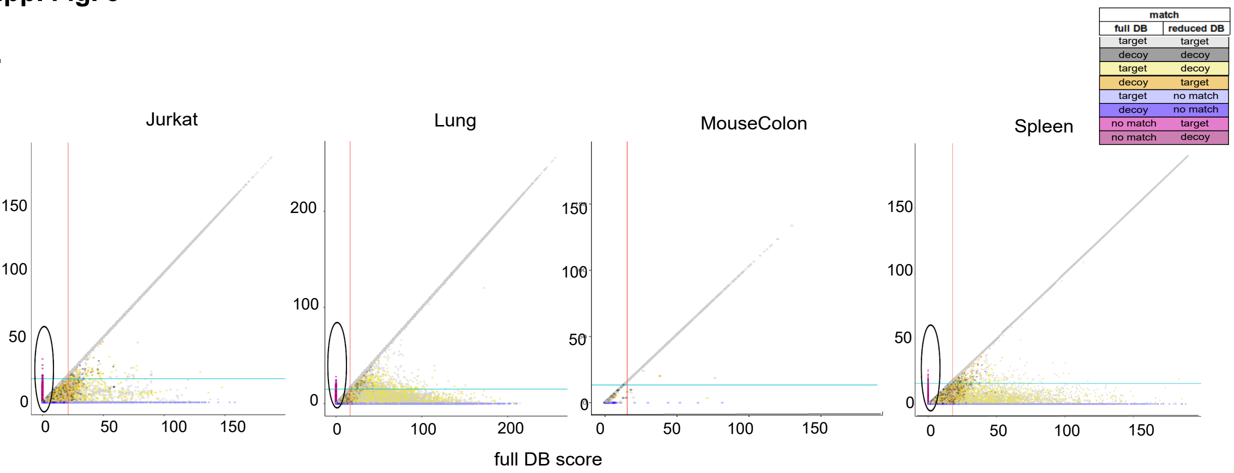


B.

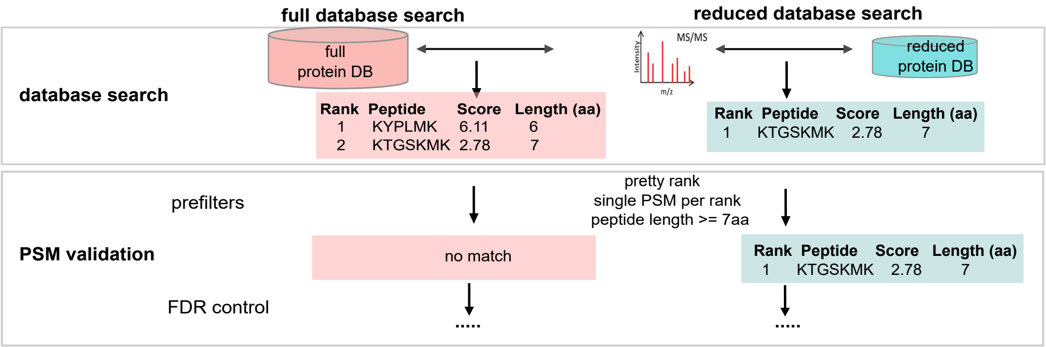


C.

| Sample | match | | nb spectra | valid at 1% FDR | score difference (reduced DB - full DB) | |
|---|---|---|---|---|---|---|
| | full DB | reduced DB | | | min | max |
| Jurkat | target | target | 49 | 1 | 0.01 | 0.1 |
| | decoy | decoy | 29 | 0 | 0.01 | 0.1 |
| | target | decoy | 56 | 1 | 0.01 | 0.1 |
| | decoy | target | 47 | 1 | 0.01 | 0.1 |
| | total | | 181 | 3 | | |

| Sample | match | | nb spectra | valid at 1% FDR | score difference (reduced DB - full DB) | |
|---|---|---|---|---|---|---|
| | full DB | reduced DB | | | min | max |
| MouseColon | target | target | 0 | 0 | - | - |
| | decoy | decoy | 0 | 0 | - | - |
| | target | decoy | 0 | 0 | - | - |
| | decoy | target | 0 | 0 | - | - |
| | total | | 0 | 0 | | |

| Sample | match | | nb spectra | valid at 1% FDR | score difference (reduced DB - full DB) | |
|---|---|---|---|---|---|---|
| | full DB | reduced DB | | | min | max |
| Lung | target | target | 226 | 4 | 0.01 | 0.1 |
| | decoy | decoy | 176 | 0 | 0.01 | 0.1 |
| | target | decoy | 235 | 4 | 0.01 | 0.09 |
| | decoy | target | 137 | 5 | 0.02 | 0.08 |
| | total | | 774 | 13 | | |

| Sample | match | | nb spectra | valid at 1% FDR | score difference (reduced DB - full DB) | |
|---|---|---|---|---|---|---|
| | full DB | reduced DB | | | min | max |
| Spleen | target | target | 287 | 15 | 0.01 | 0.01 |
| | decoy | decoy | 271 | 1 | 0.01 | 0.01 |
| | target | decoy | 282 | 1 | 0.01 | 0.01 |
| | decoy | target | 205 | 0 | 0.01 | 0.01 |
| | total | | 1045 | 17 | | |

D.

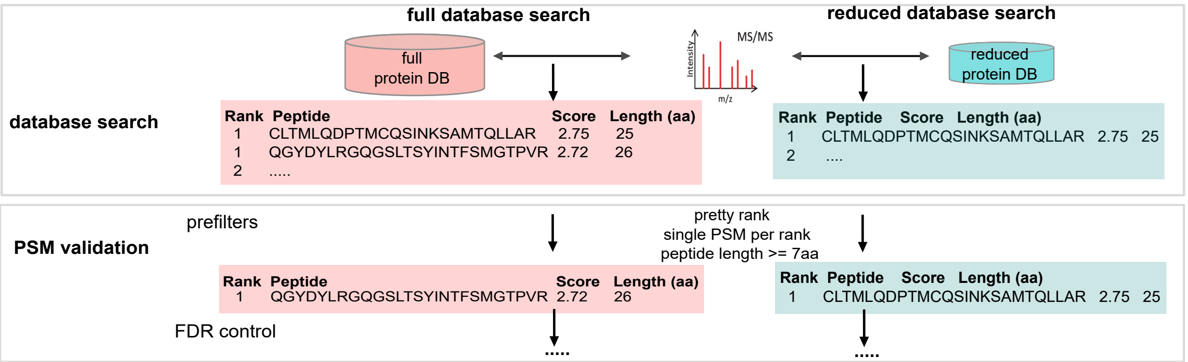

**Supp. Fig. 4**
Jurkat 1% FDR
(TDC)

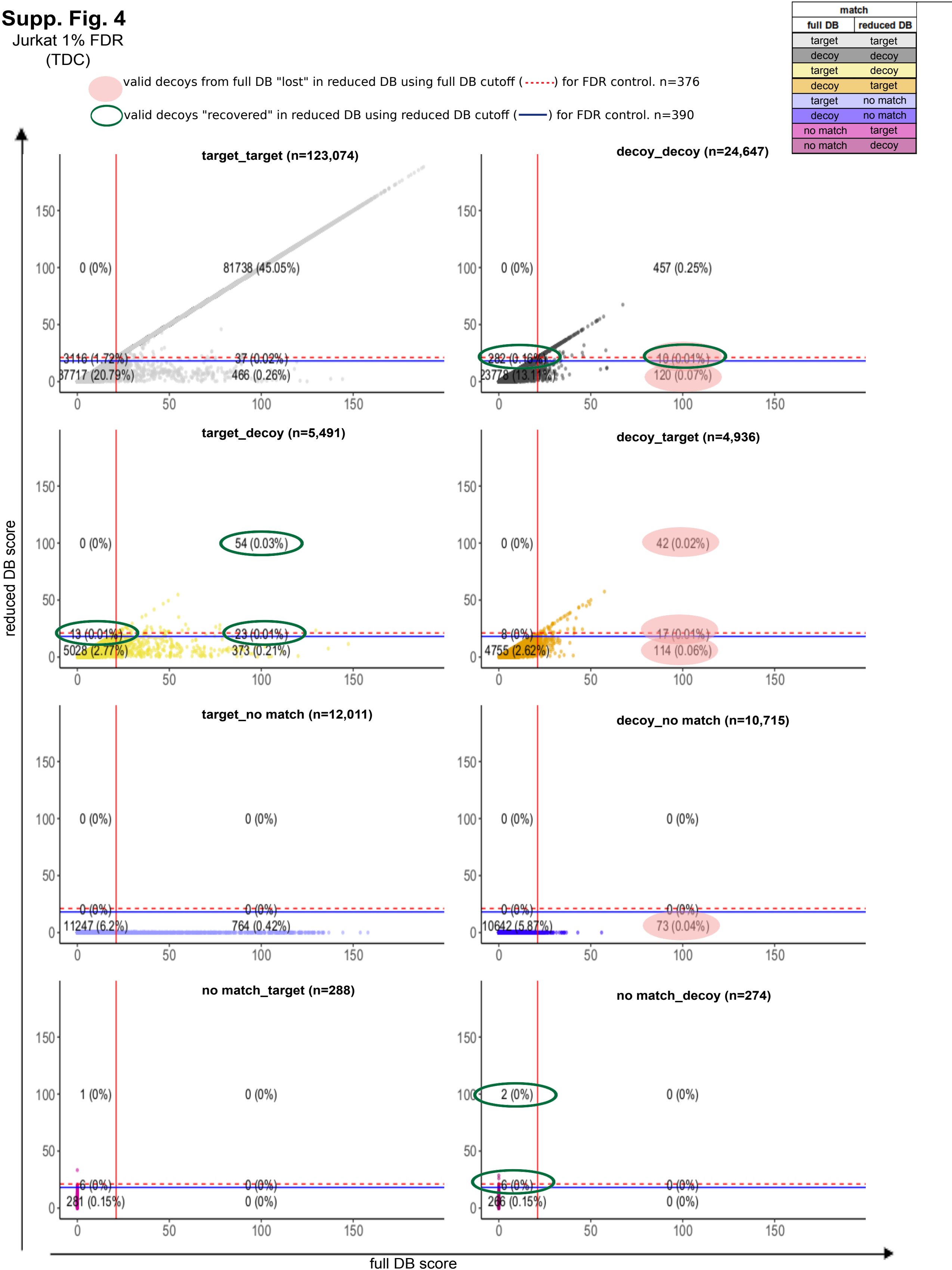

**Supp. Fig. 5**

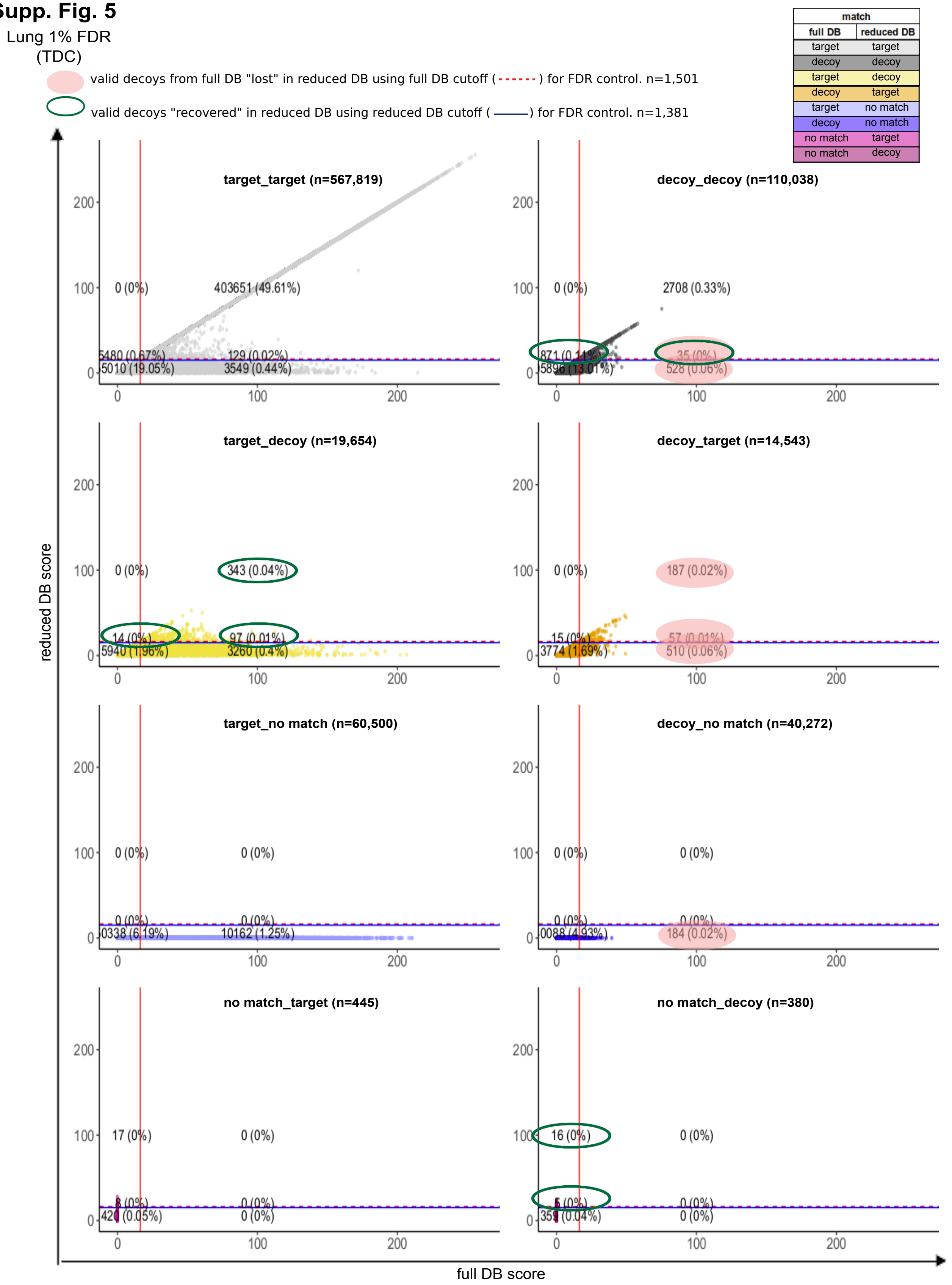

Lung 1% FDR
(TDC)
valid decoys from full DB "lost" in reduced DB using full DB cutoff ( ) for FDR control. n=1,501
valid decoys "recovered" in reduced DB using reduced DB cutoff ( ) for FDR control. n=1,381
match
full DB
reduced DB
target target
decoy decoy
target decoy
decoy target
target no match
decoy no match
no match target
no match decoy
target_target (n=567,819)
0 (0%)
403651 (49.61%)
5480 (0.67%)
129 (0.02%)
5010 (19.05%)
3549 (0.44%)
decoy_decoy (n=110,038)
0 (0%)
2708 (0.33%)
871 (0.11%)
35 (0%)
5896 (13.01%)
528 (0.06%)
target_decoy (n=19,654)
0 (0%)
343 (0.04%)
14 (0%)
97 (0.01%)
5940 (1.96%)
3260 (0.4%)
decoy_target (n=14,543)
0 (0%)
187 (0.02%)
15 (0%)
57 (0.01%)
3774 (1.69%)
510 (0.06%)
target_no match (n=60,500)
0 (0%)
0 (0%)
0 (0%)
0 (0%)
0338 (6.19%)
10162 (1.25%)
decoy_no match (n=40,272)
0 (0%)
0 (0%)
0 (0%)
0 (0%)
0088 (4.93%)
184 (0.02%)
no match_target (n=445)
17 (0%)
0 (0%)
8 (0%)
0 (0%)
420 (0.05%)
0 (0%)
no match_decoy (n=380)
16 (0%)
0 (0%)
5 (0%)
0 (0%)
359 (0.04%)
0 (0%)
reduced DB score
full DB score

**Supp. Fig. 6**

MouseColon 1% FDR
(TDC)

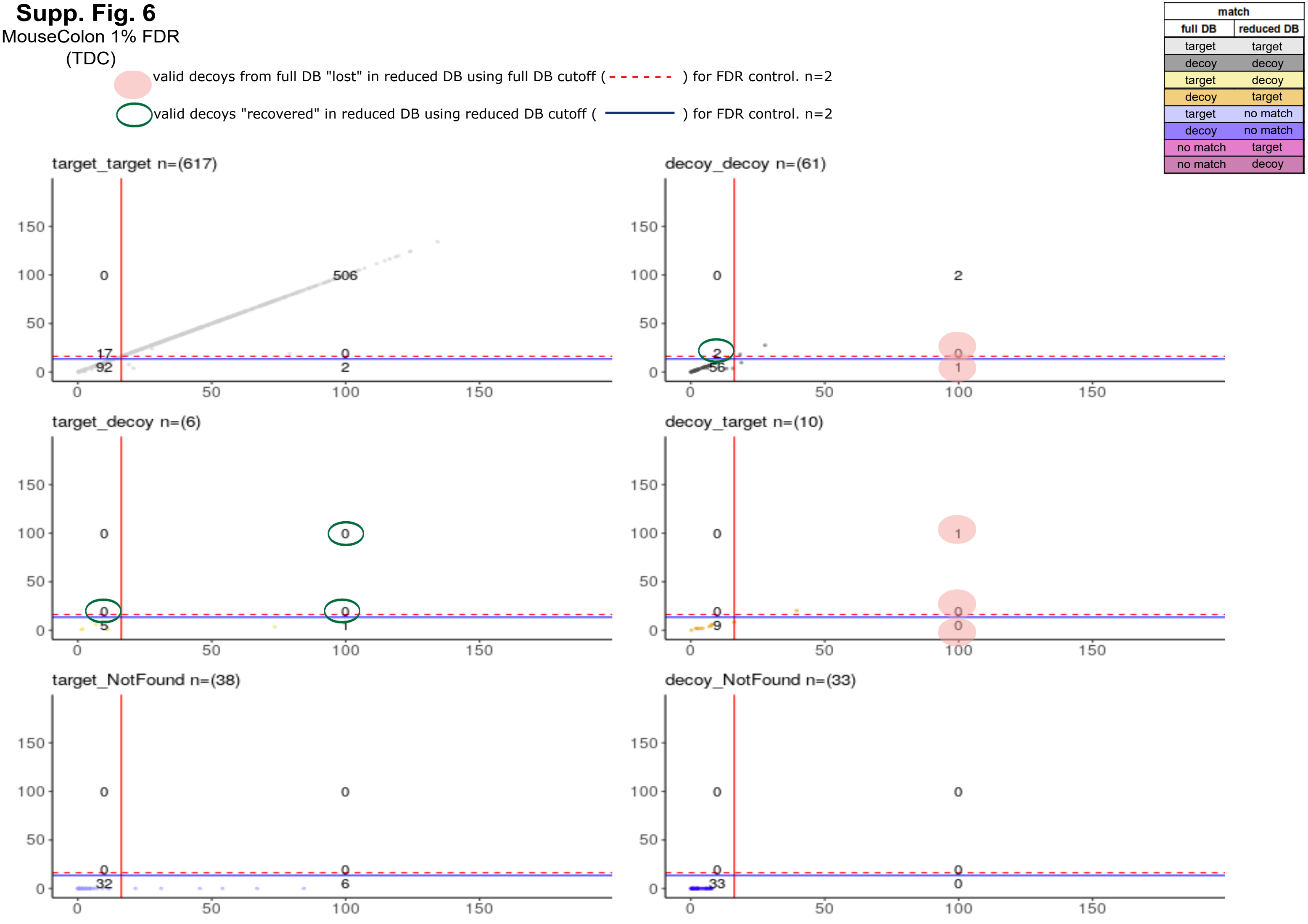

### Supp. Fig. 7

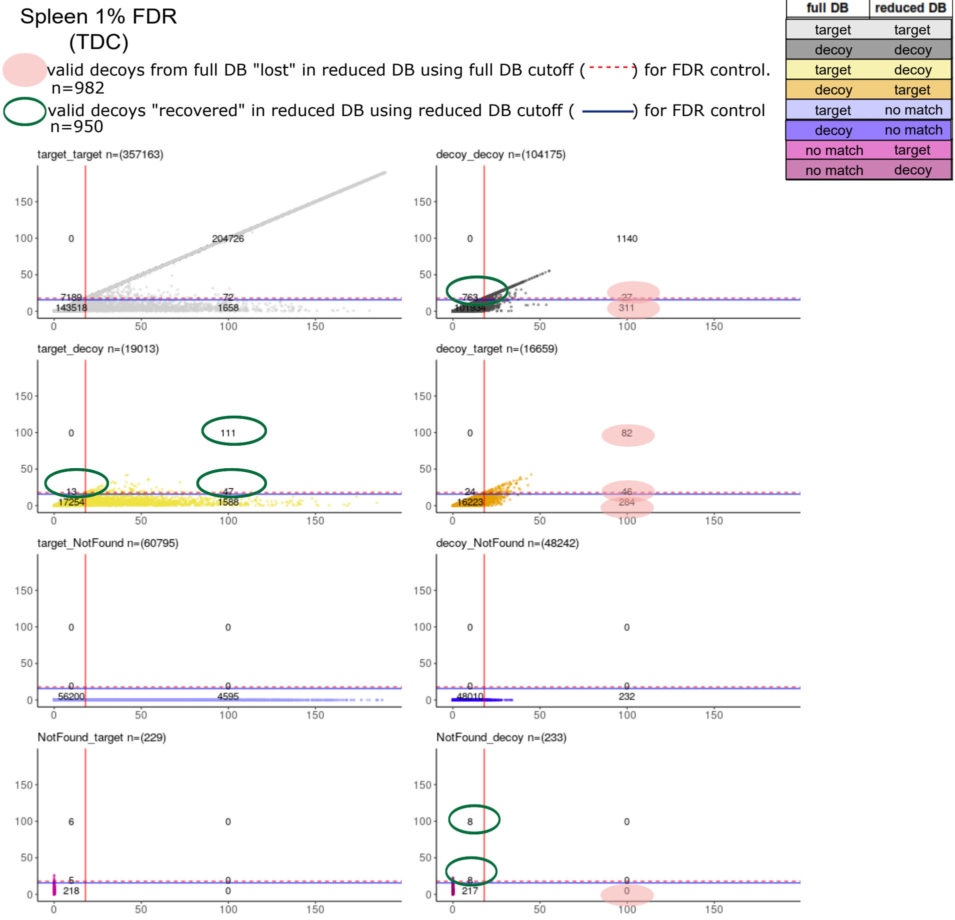
Spleen 1% FDR
(TDC)
valid decoys from full DB "lost" in reduced DB using full DB cutoff ( ) for FDR control.
n=982
valid decoys "recovered" in reduced DB using reduced DB cutoff ( ) for FDR control
n=950
match
full DB
reduced DB
target target
decoy decoy
target decoy
decoy target
target no match
decoy no match
no match target
no match decoy
target_target n=(357163)
0
204726
7189
72
143518
1658
decoy_decoy n=(104175)
0
1140
763
27
101934
311
target_decoy n=(19013)
0
111
13
47
17254
1588
decoy_target n=(16659)
0
82
24
46
16223
284
target_NotFound n=(60795)
0
0
0
0
56200
4595
decoy_NotFound n=(48242)
0
0
0
0
48010
232
NotFound_target n=(229)
6
0
5
0
218
0
NotFound_decoy n=(233)
8
0
8
0
217
0
0
50
100
150

### Supp. Fig. 8

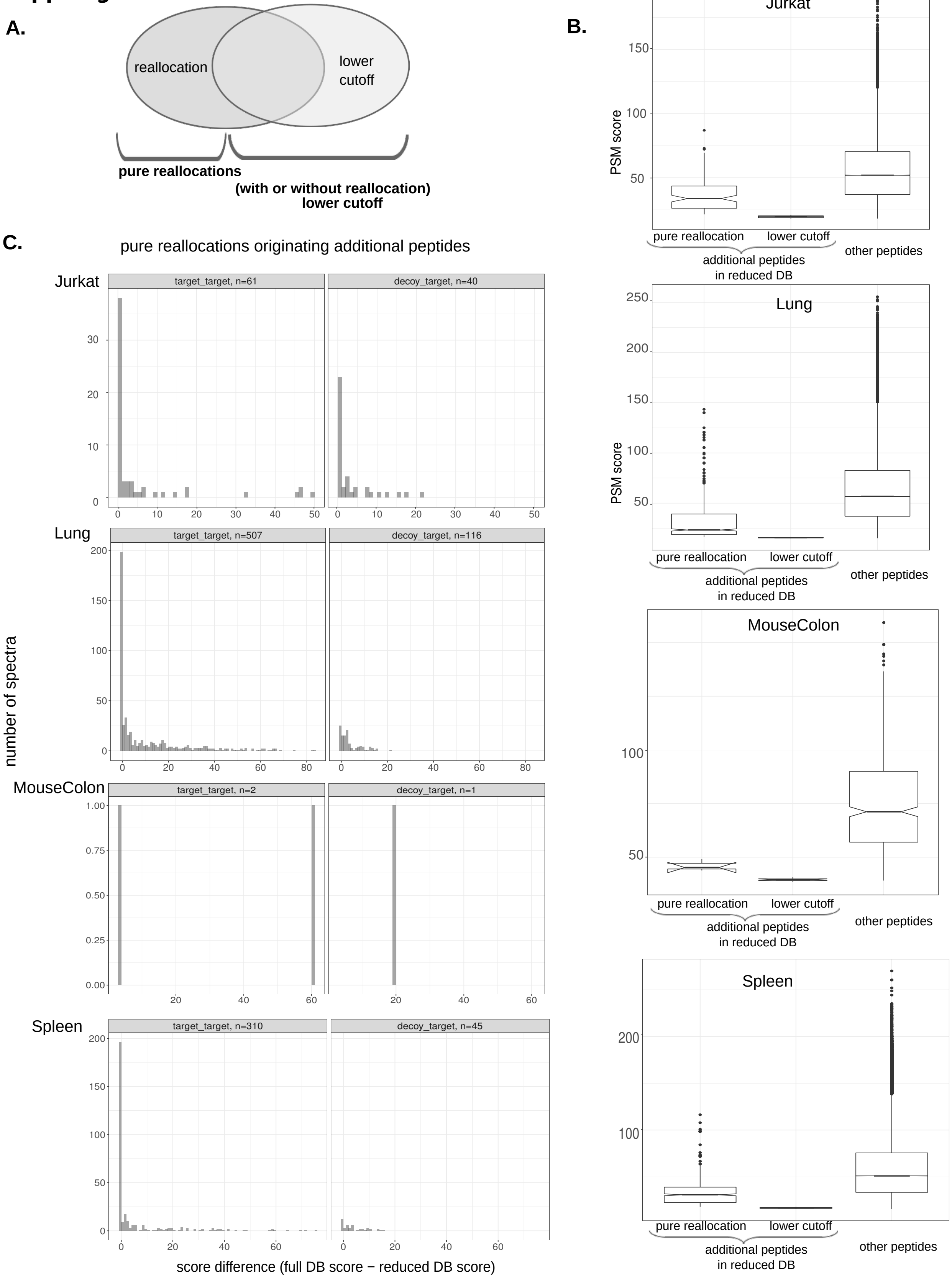
A.
reallocation
lower
cutoff
pure reallocations
(with or without reallocation)
lower cutoff
B.
Jurkat
PSM score
pure reallocation
lower cutoff
additional peptides
in reduced DB
other peptides
Lung
MouseColon
Spleen
C.
pure reallocations originating additional peptides
Jurkat
target_target, n=61
decoy_target, n=40
Lung
target_target, n=507
decoy_target, n=116
MouseColon
target_target, n=2
decoy_target, n=1
Spleen
target_target, n=310
decoy_target, n=45
number of spectra
score difference (full DB score − reduced DB score)

**Supp. Fig. 9**

**A.**

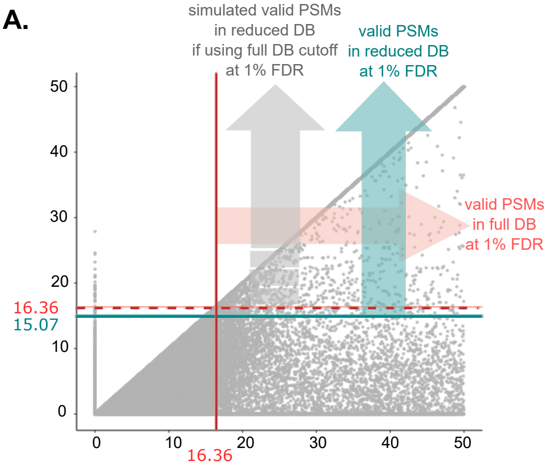


**B.**

| Database | Score cutoff | # valid targets | # valid decoys | FDR |
|---|---|---|---|---|
| full | 16.36 | 421,191 | 4,209 | 1 |
| reduced | 16.36 | 403,855 | 3,067 | 0.76 |
| reduced | 15.07 | 409,544 | 4,089 | 1 |

-4.1% -27.1%

+1.3% +24.3%

**C.**

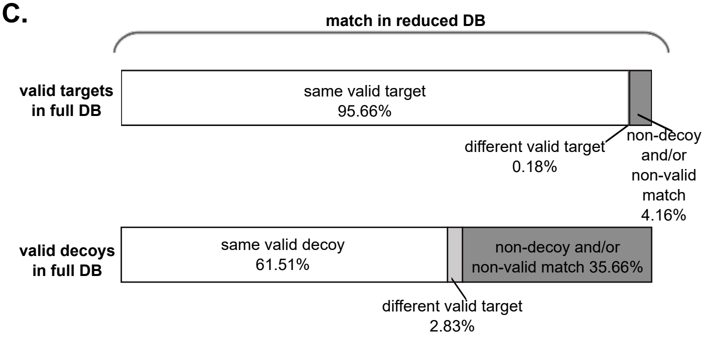


**D.**

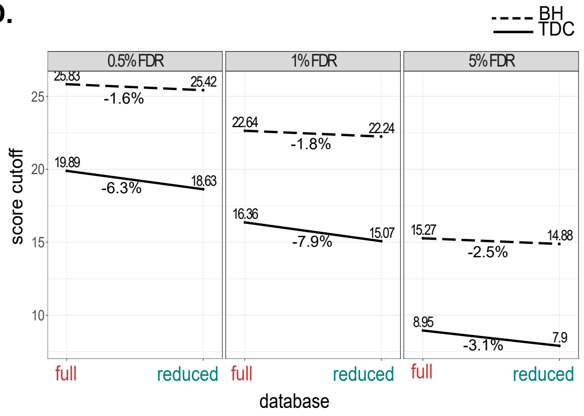

**Supp. Fig. 10**

A.

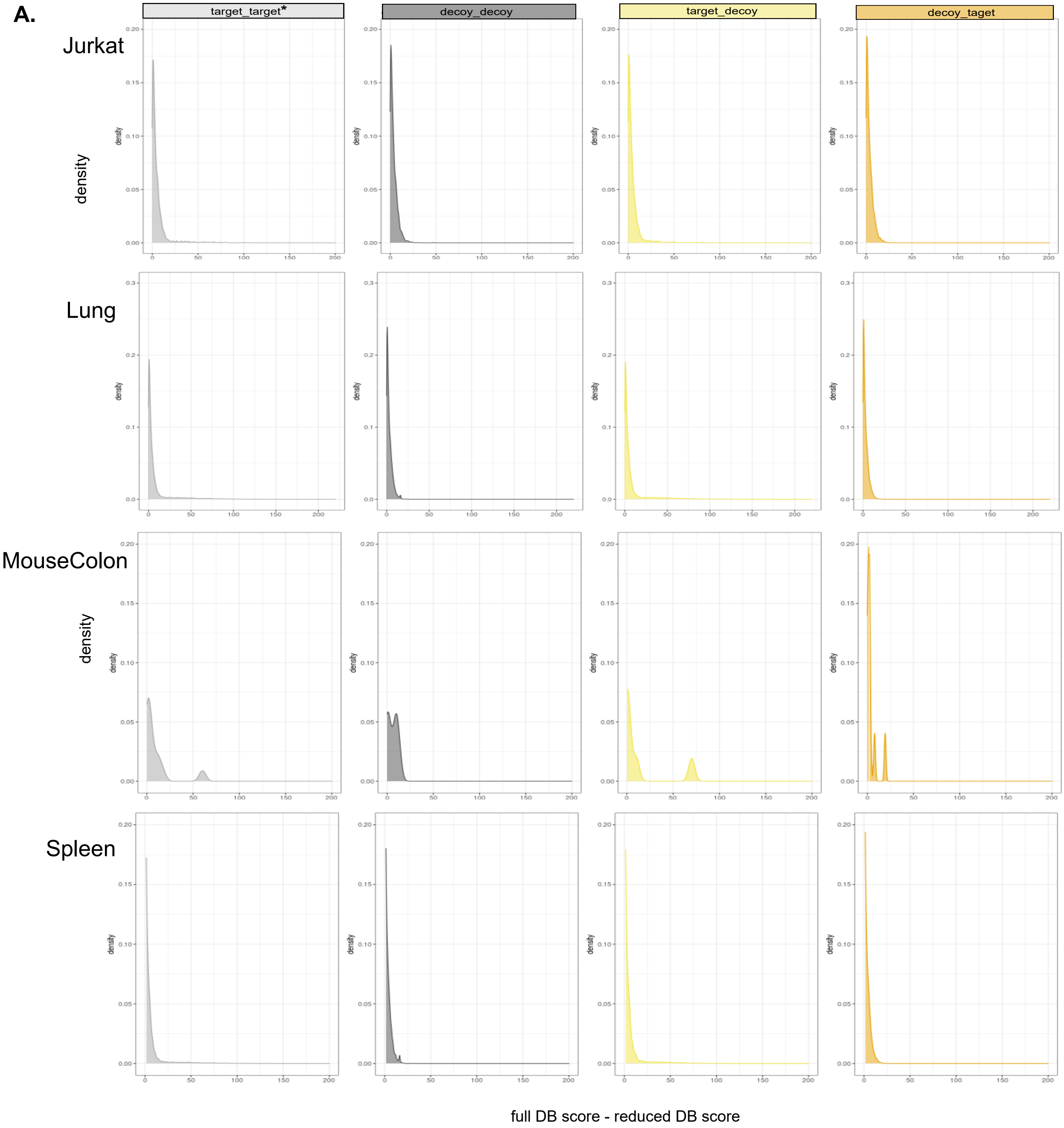


B.

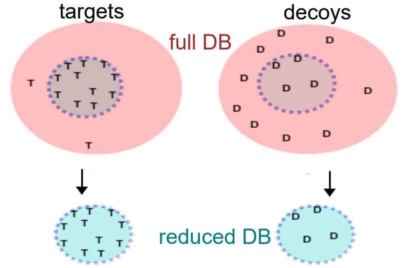

#### Supp. Fig. 11

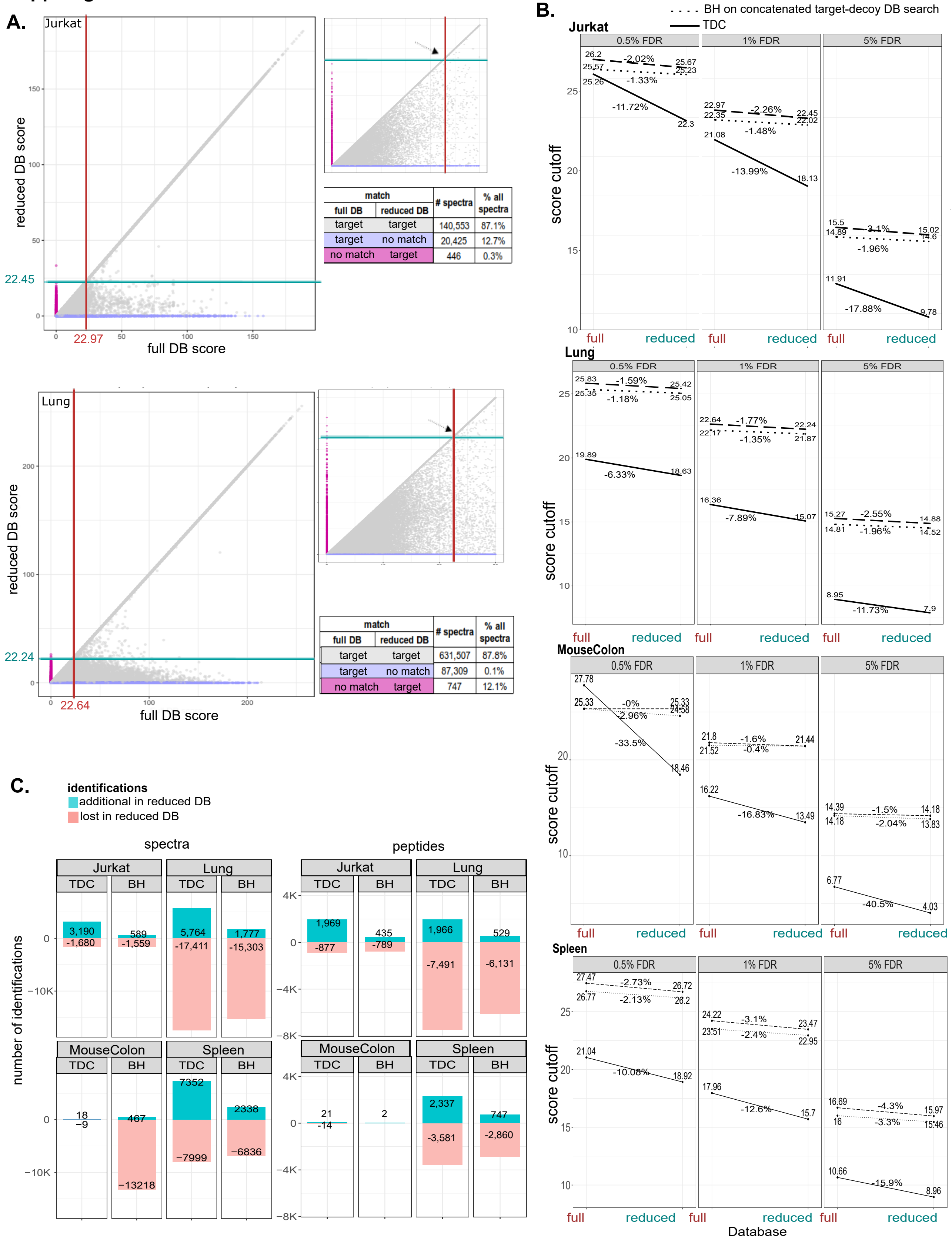


Jurkat:

| match | | # spectra | % all spectra |
|---|---|---|---|
| full DB | reduced DB | | |
| target | target | 140,553 | 87.1% |
| target | no match | 20,425 | 12.7% |
| no match | target | 446 | 0.3% |

Lung:

| match | | # spectra | % all spectra |
|---|---|---|---|
| full DB | reduced DB | | |
| target | target | 631,507 | 87.8% |
| target | no match | 87,309 | 0.1% |
| no match | target | 747 | 12.1% |

### Supp Fig 12

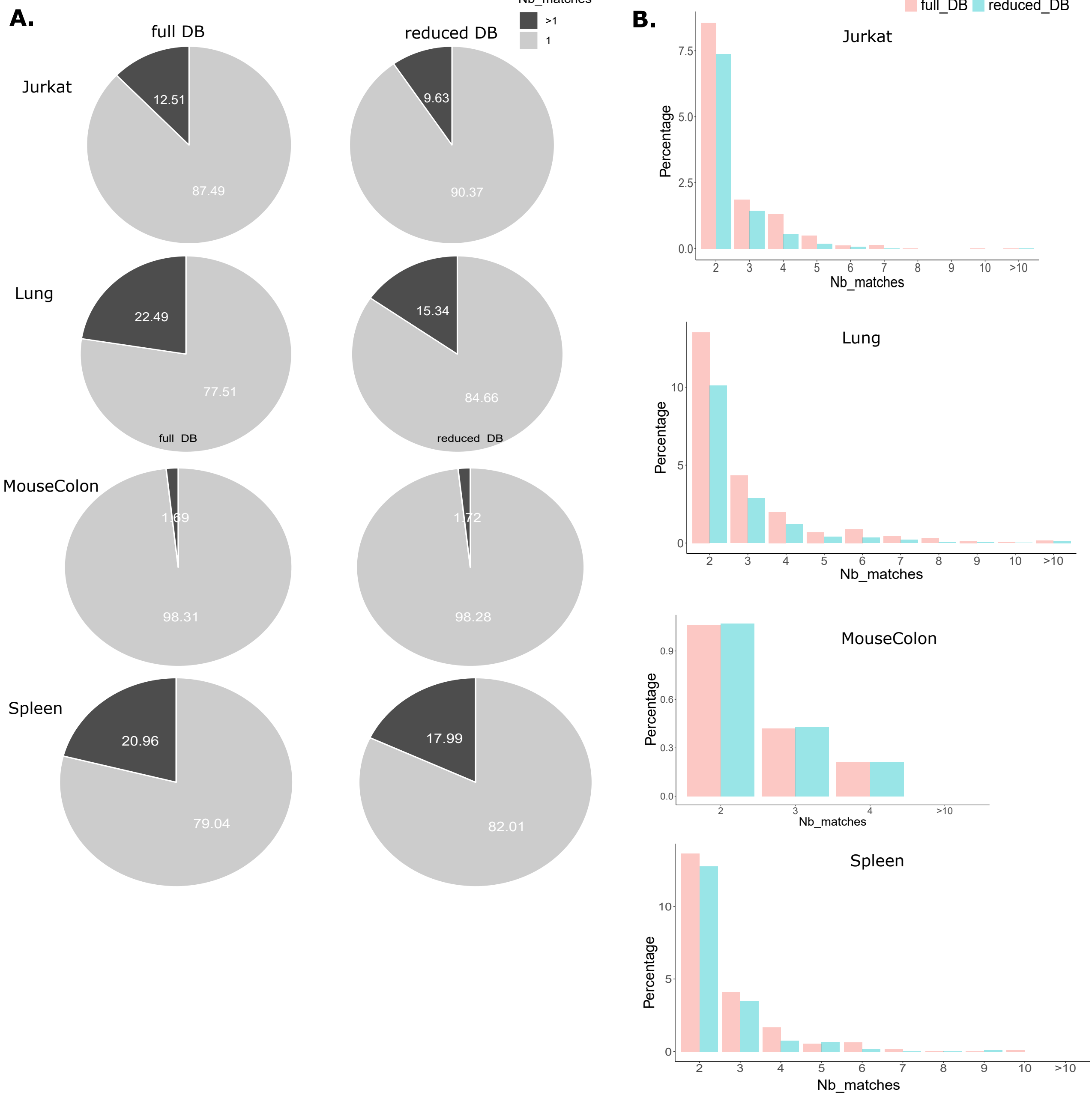

A.
full DB
reduced DB
Nb_matches
>1
1
Jurkat
12.51
87.49
9.63
90.37
Lung
22.49
77.51
15.34
84.66
full DB
reduced DB
MouseColon
1.69
98.31
1.72
98.28
Spleen
20.96
79.04
17.99
82.01
B.
full_DB
reduced_DB
Jurkat
Lung
MouseColon
Spleen
Percentage
Nb_matches

**Supp. Fig. 13**

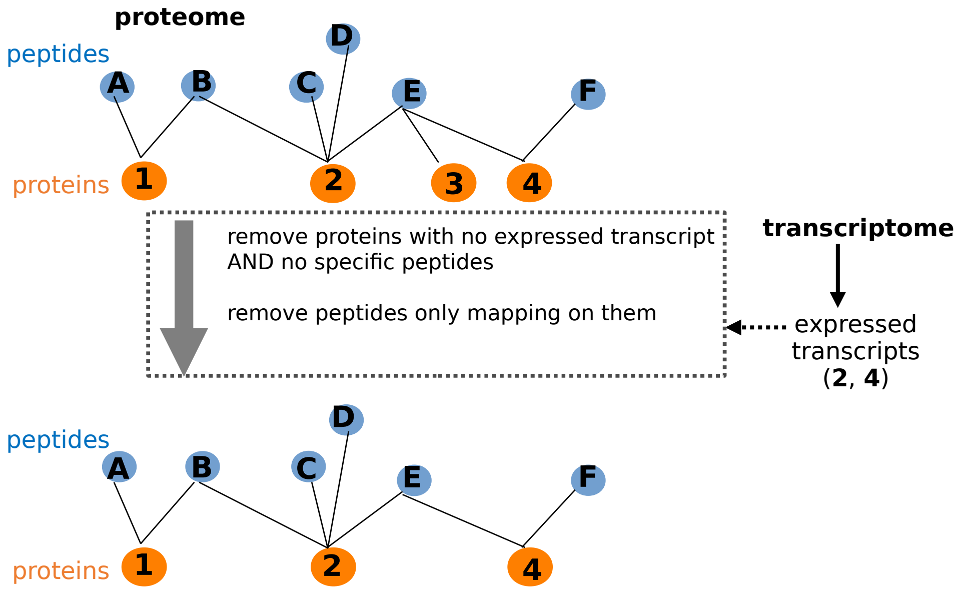
proteome
peptides
A
B
C
D
E
F
proteins
1
2
3
4
remove proteins with no expressed transcript AND no specific peptides
remove peptides only mapping on them
transcriptome
expressed transcripts (2, 4)
peptides
A
B
C
D
E
F
proteins
1
2
4

**Supp. Fig. 14**

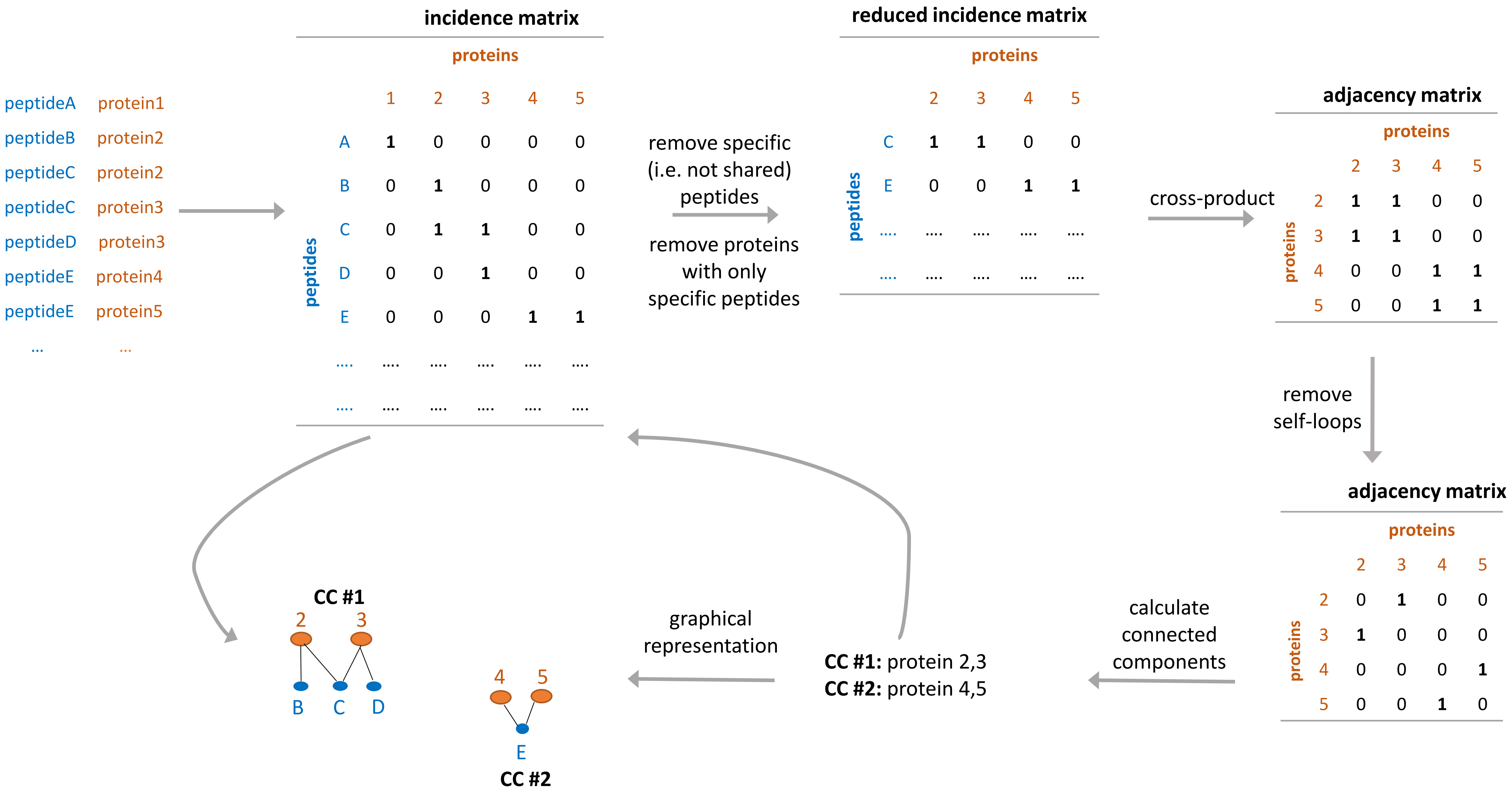
peptideA protein1
peptideB protein2
peptideC protein2
peptideC protein3
peptideD protein3
peptideE protein4
peptideE protein5
... ...
incidence matrix
proteins
1 2 3 4 5
peptides
A 1 0 0 0 0
B 0 1 0 0 0
C 0 1 1 0 0
D 0 0 1 0 0
E 0 0 0 1 1
.... .... .... .... .... ....
.... .... .... .... .... ....
remove specific (i.e. not shared) peptides
remove proteins with only specific peptides
reduced incidence matrix
proteins
2 3 4 5
peptides
C 1 1 0 0
E 0 0 1 1
.... .... .... .... ....
.... .... .... .... ....
cross-product
adjacency matrix
proteins
2 3 4 5
2 1 1 0 0
3 1 1 0 0
4 0 0 1 1
5 0 0 1 1
remove self-loops
adjacency matrix
proteins
2 3 4 5
2 0 1 0 0
3 1 0 0 0
4 0 0 0 1
5 0 0 1 0
calculate connected components
CC #1: protein 2,3
CC #2: protein 4,5
graphical representation
CC #1
2 3
B C D
4 5
E
CC #2
