## Additional File 3 for "Transcriptome-informed reduction of protein databases: an analysis of how and when proteogenomics enhances eukaryotic proteomics"

### SUPPLEMENTARY NOTES

#### Supplementary Note 1: PSM score comparison between full and reduced database searches

We carefully compared all (target or decoy) PSMs passing validation prefilters (pretty rank; single PSM per rank; minimum peptide length of 7 amino acids) from the full and reduced database search, prior to filtering for 1% FDR control and we systematically verified that in no case a spectrum has a match with higher score in the reduced database than in the full database search. This due to the fact that the Mascot score for equivalent PSMs is the same independently of the database searched, as already described (24), and, since the reduced database is a strict subset of the full database, PSMs from the former can only present lower or at best equal scores than the latter. We verified all those cases which may appear to contradict this assumption. First, we examined spectra with a match passing prefilters only in the reduced database (indicated as “no match, target” or “no match, decoy” in **Figure 2A, Supp. Fig. 2A, Supp. Fig. 3A, Supp. Fig. 4 and 5**). To be able to represent them in a plot, we arbitrarily assigned them a score of zero in the full database, while they actually have no score at all in the full database: they just have no valid match. The absence of valid match in the full database, while present in the reduced database, is explained by reallocation and prefilters used for validation. Their best PSM in the full database is discarded as it does not satisfy the minimum peptide length prefilter (>= 7 amino acids). On the contrary, in the reduced database, since the peptide match from the full database is not present, they are reallocated to a longer peptide, which passes prefilters so that the match is retained (**Supp. Fig. 3B**). Therefore, these spectra do not contradict the assumption that PSMs from the reduced database search can only present lower or at best equal scores than in the full database. Next, we examined the 181 spectra (0.1% of all spectra, including 2 target matches validated at 1% FDR in reduced database) in Jurkat and 774 spectra (0.1% of all spectra, including 9 target matches validated at 1% FDR in reduced database) in Lung (**Supp. Fig. 3C, Supp. Fig 4 and 5**), with a score difference (of 0.1 at best) in favor of the reduced database. These spectra do not really present a higher PSM score in the reduced database but they are explained by the use of the “pretty rank” prefilter (**Supp. Fig. 3D**). Indeed, by using “pretty rank” we consider of equal score all PSMs for the same spectrum with a score difference inferior to 0.1 and assign them the same rank, the reason being that a score difference of 0.1 is most likely too small to rank PSMs accordingly. In some cases, this results in multiple PSMs of rank 1, with a slight difference in score (up to 0.1). We retained a single best PSM per spectrum (“single PSM per rank” filter) to make the analysis more interpretable and, in some cases, the rank 1 PSM with the lower score was retained for the reduced database[1]. All above described spectra present at least one PSM in the full database search whose score is higher or equal to that from the reduced database, but not retained after single PSM per rank filter. This confirms again that in no case the score from the reduced database is higher than in the full database.

*[1] In case of several PSMs with the same rank, the “single PSM per rank” filter implemented in the Proline software chooses the PSM identifying the protein with the maximum number of valid PSMs; if equality, it chooses the PSM with the lower delta m/z and if still equality, it randomly selects one.*

### Supplementary Note 2: Transcriptome information helps to reduce ambiguity at the PSM level

We observed that searches against reduced databases were associated with slightly lower ambiguity at the PSM level, in addition to lower ambiguity at the protein level. In the spectrum identification step, it is common to only consider the best peptide match for each spectrum (*i.e.*, the rank 1 PSM, according to the search engine score) but a spectrum may match different peptides equally (or almost equally) well. This complicates the analysis, and no consensus exists on how to treat these cases. Interestingly, we observed that a smaller proportion of spectra were associated with multiple best matches in the reduced database search (**Supp. Fig. 10A**); likewise, fewer best matches were generally found per spectrum (**Supp. Fig. 10B**).
